# Integrated omics analysis unveils a DNA damage response to neurogenic injury

**DOI:** 10.1101/2023.12.10.571015

**Authors:** Ali Hashemi Gheinani, Bryan S. Sack, Alex Bigger-Allen, Hatim Thaker, Hussein Atta, George Lambrinos, Kyle Costa, Claire Doyle, Mehrnaz Gharaee-Kermani, Susan Patalano, Mary Piper, Justin F. Cotellessa, Dijana Vitko, Haiying Li, Manubhai Kadayil Prabhakaran, Vivian Cristofaro, John Froehlich, Richard S. Lee, Wei Yang, Maryrose P. Sullivan, Jill A. Macoska, Rosalyn M. Adam

**Affiliations:** Urological Diseases Research Center, Boston Children’s Hospital, Boston, MA, USA; Functional Urology Research Group, Department for BioMedical Research DBMR, University of Bern, Switzerland; Department of Urology, Inselspital University Hospital, 3010 Bern, Switzerland; Department of Surgery, Harvard Medical School, Boston, MA, USA; Broad Institute of MIT and Harvard, Cambridge, MA, USA; Biological & Biomedical Sciences Graduate Program, Division of Medical Sciences, Harvard Medical School, Boston, MA; University of Massachusetts, Boston, MA, USA; Harvard Chan Bioinformatics Core, Harvard T.H. Chan School of Public Health, Boston, MA, USA; Division of Urology, VA Boston Healthcare System, Boston, MA, USA; Departments of Surgery and Biomedical Sciences, Cedars-Sinai Medical Center, Los Angeles, CA

**Keywords:** PARP-1 activation, DNA damage, tissue remodeling, neurogenic bladder

## Abstract

Spinal cord injury (SCI) evokes profound bladder dysfunction. Current treatments are limited by a lack of molecular data to inform novel therapeutic avenues. Previously, we showed systemic inosine treatment improved bladder function following SCI in rats. Here, we applied multi-omics analysis to explore molecular alterations in the bladder and their sensitivity to inosine following SCI. Canonical pathways regulated by SCI included those associated with protein synthesis, neuroplasticity, wound healing, and neurotransmitter degradation. Upstream regulator analysis identified MYC as a key regulator, whereas causal network analysis predicted multiple regulators of DNA damage response signaling following injury, including PARP-1. Staining for both DNA damage (γH2AX) and PARP activity (poly-ADP-ribose) markers in the bladder was increased following SCI, and attenuated in inosine-treated tissues. Proteomics analysis suggested that SCI induced changes in protein synthesis-, neuroplasticity-, and oxidative stress-associated pathways, a subset of which were shown in transcriptomics data to be inosine-sensitive. These findings provide novel insights into the molecular landscape of the bladder following SCI, and highlight a potential role for PARP inhibition to treat neurogenic bladder dysfunction.

**Synopsis:** 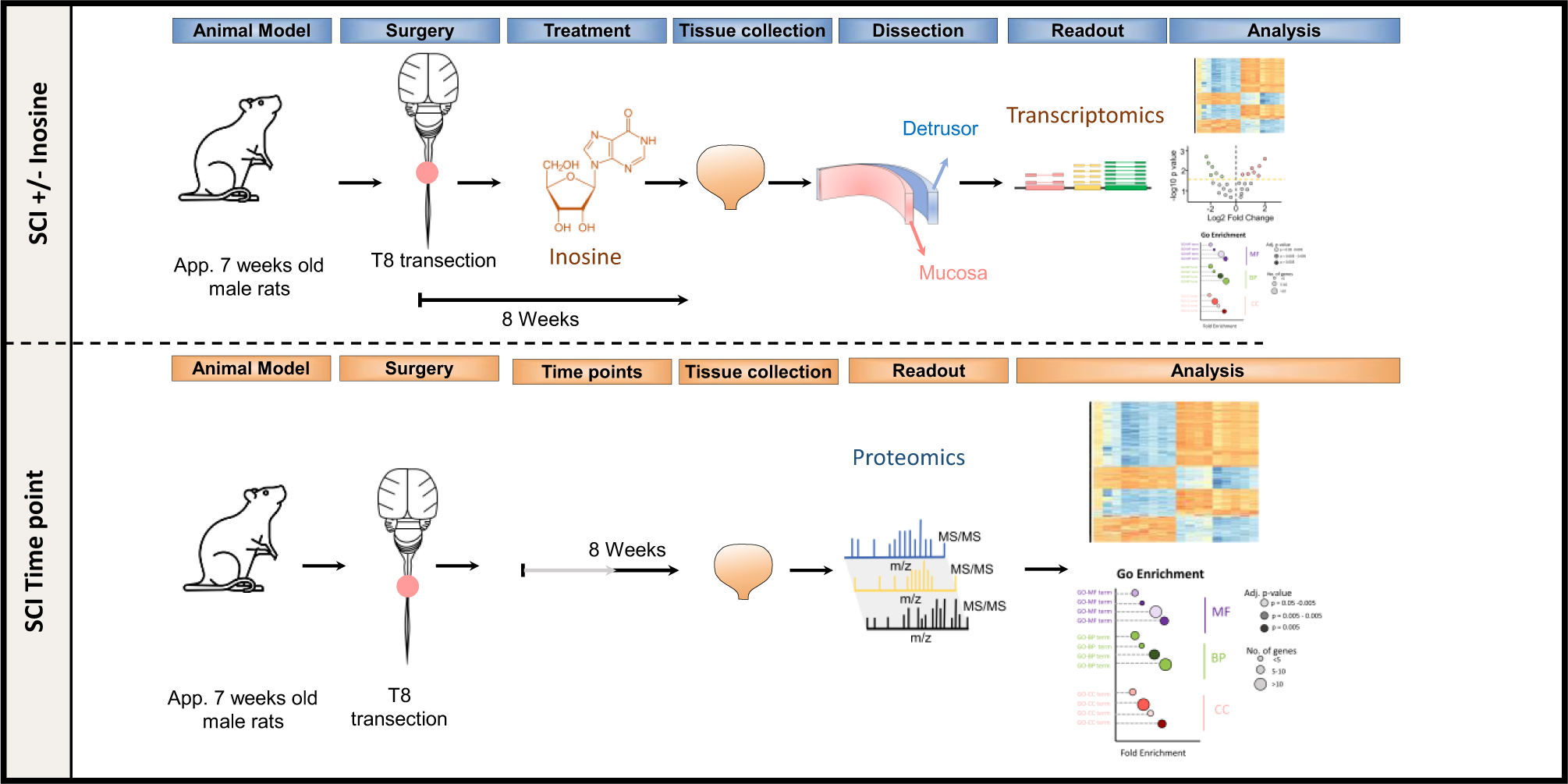

- Employed a multi-omics approach, integrating both transcriptomic and proteomic analyses, to investigate the molecular response in a rat model of spinal cord injury (SCI) and the therapeutic effect of inosine.
- Discovered multiple regulators of the DNA damage response, including PARP-1, using causal network analysis.
- Observed decreased markers of DNA damage and PARP activity in inosine-treated tissues, indicating the therapeutic potential of inosine in neurogenic dysfunction.
- Identified significant alterations in molecular pathways associated with protein synthesis, neuroplasticity, wound healing, and neurotransmitter degradation after SCI, and their modulation by inosine, highlighting its neuroprotective effects beyond DNA damage repair.

## Introduction

Damage to the spinal cord has profound impact on motor, sensory and autonomic innervation. Depending on the level and extent, injury can result in limb paralysis, bladder/bowel dysfunction and development of neuropathic pain (reviewed in [1]). Interruption of neural control to the lower urinary tract frequently leads to reflex bladder contractions termed neurogenic detrusor overactivity (NDO) and detrusor-sphincter-dyssynergia (DSD) that reflects disruption of the normal coordination between bladder and urethral sphincter required for efficient voiding. NDO and DSD are associated with emergence of lower urinary tract symptoms including urinary incontinence, incomplete bladder emptying resulting in urinary retention, and an increased risk of urinary tract infection and renal damage (reviewed in [2]). Clinical management of neurogenic lower urinary tract dysfunction (LUTD) focuses primarily on catheterization to promote efficient bladder emptying, and pharmacological intervention to diminish NDO. Agents approved for treatment of LUTD include onabotulinumtoxin A to inhibit neurotransmitter release, antimuscarinic agents to block muscarinic receptor-mediated detrusor contraction, and β-adrenergic receptor agonists to promote detrusor relaxation [3]. These interventions decrease bladder storage pressure, enhance bladder capacity and diminish incontinence.

The functional obstruction that results from DSD also provokes marked changes in bladder morphology, consistent with those observed following anatomic obstruction e.g. in response to prostatic enlargement in men. Characteristic changes include bladder wall thickening as a result of cellular hypertrophy and hyperplasia [4], as well as enhanced deposition and turnover of extracellular matrix (ECM) leading to fibrosis [5, 6]. With the advent of genome-wide expression profiling, several groups have begun to characterize the molecular landscape that underlies the response of the bladder to spinal cord damage in animal models [7–10] and in smooth muscle cells isolated from human bladder biopsies [11, 12]. In addition to verification of altered expression of ECM proteins and regulators, microarray-based expression profiling of bladder tissue from spinal cord injured rats also provided important insights into additional drivers of neurogenic bladder pathophysiology including the pro-fibrotic regulator TGFβ1 and pro-inflammatory mediators such as IL1β and S100A8/A9 [7, 8]. In those studies, pathway analysis identified the enrichment for processes associated with remodeling such as proliferation and connective tissue deposition, as well as a progressive increase in inflammatory processes following injury. In spite of its deleterious impact on bladder compliance and function, however, no pharmacological agents that target fibroproliferative remodeling have been approved to treat LUTD. While these studies have provided valuable insights into molecular alterations associated with spinal cord injury (SCI), the lack of high-resolution, multi-dimensional and network-level information has hindered the development of rationally-designed therapeutic interventions targeting bladder wall remodeling.

Previously, we demonstrated a significant improvement in bladder function in rats with functional bladder outlet obstruction secondary to SCI treated with inosine [13], a purine nucleoside with neuroprotective, neurotrophic, anti-inflammatory and anti-oxidant properties (reviewed in [14]). In that study, rats with SCI receiving inosine daily for 6 weeks either immediately after injury or following a two-month delay displayed a decrease in NDO. Consistent with the neuroprotective activity of inosine reported previously, we observed preservation of the neuronal markers synaptophysin and NF200 within the bladders of inosine-treated rats, as well as decreased staining for TRPV1, a marker of C-fibers implicated in the development of NDO following injury (reviewed in [15]).

To determine the molecular basis for the beneficial impact of inosine on bladder function observed previously, we conducted a multi-omics analysis to systematically evaluate the genes, pathways and signaling networks that are differentially regulated in the bladder in response to SCI and their modulation with inosine treatment. This multi-omics analysis has provided novel insights into the biological processes perturbed in the bladder in response to SCI, and identified PARP-1 as a novel effector of inosine.

## Results

### Inosine prevents SCI-induced transcriptome changes in detrusor and mucosa

Previous studies from our group have shown that chronic inosine treatment can mitigate neurogenic detrusor overactivity in spinal cord-injured rats independently of direct effects on muscle contractility [13, 16] (Suppl. Fig. 1A-C). Whereas SCI was associated with decreased levels of synaptophysin and the Aδ fiber marker NF200 in the bladder wall, compared to non-injured controls, inosine treatment preserved synaptophysin and NF200 levels, consistent with its known neuroprotective activity [14] (Suppl. Fig. 1A-C). To further interrogate the impact of inosine in the bladder following SCI we performed transcriptomics analysis on bladder tissue from rats subjected to complete mid-thoracic spinal cord transection and treated with inosine or vehicle for 8 weeks after injury (Fig. 1A). Consistent with previous findings from us and others [17–19], the bladder-to-body weight ratios in SCI rats were significantly higher than in control animals, primarily as a result of increased bladder weights following injury (*, p < 0.05 in each case, Figure 1B). RNA was isolated from representative detrusor and mucosa samples microdissected from bladders at the time of harvest (Fig. 1A) and subjected to RNA sequencing (RNA-seq). Profiling captured differentially expressed RNAs in different groups (Suppl. Fig. 1A-C).

**Figure 1.**
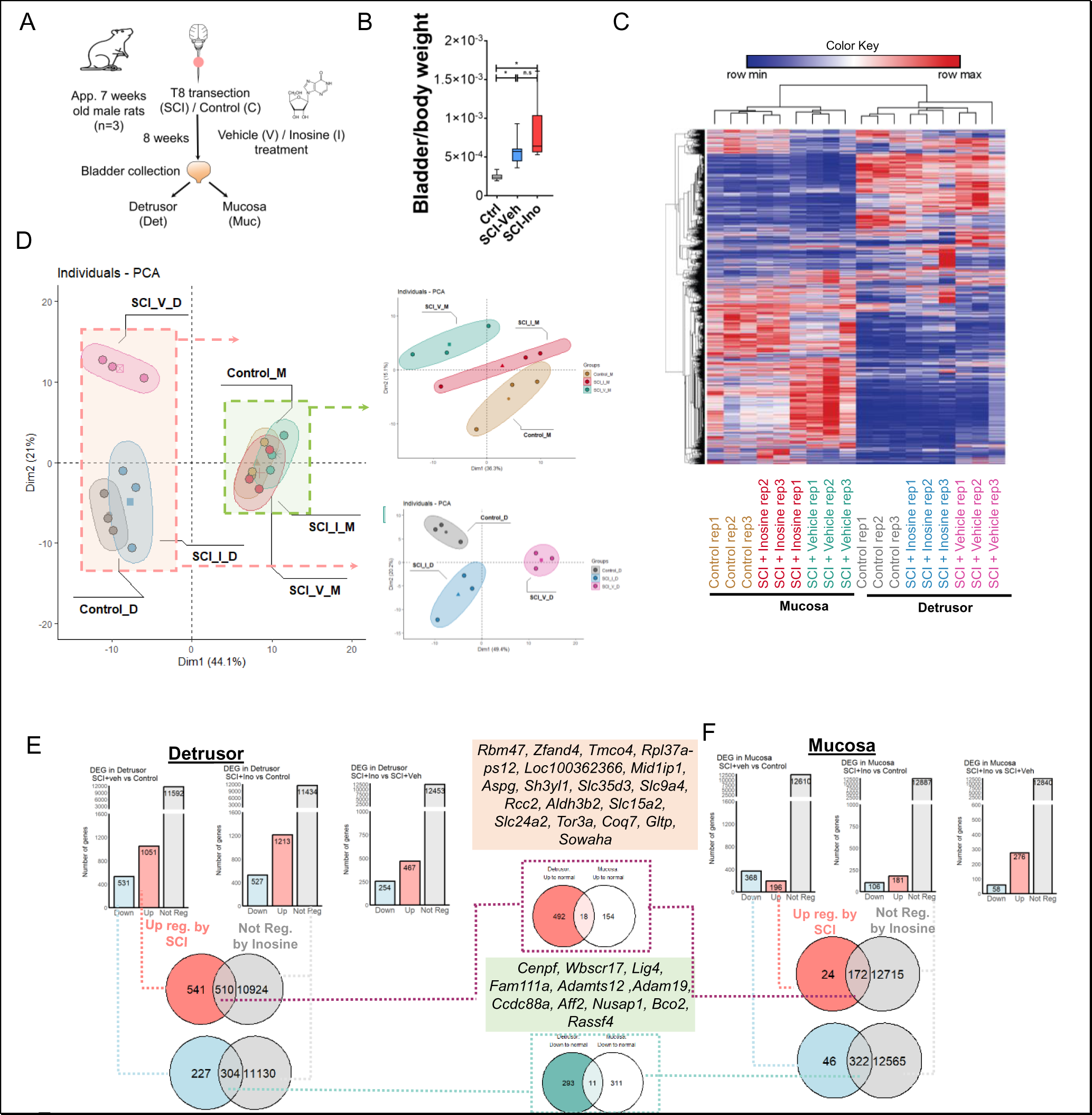
Effect of spinal cord injury and inosine treatment on transcriptome of bladder wall. (A) Experimental Design. (B) Bladder-to-body weight ratio measurement. (C) Heatmap and hierarchical clustering using DEGs in different comparisons groups. log2 fold change> +/- 0.5; P-value <0.05; read counts cpm>1 (D) PCA based on top 200 variable genes. (E) Bar charts indicating upregulated and down regulated DEGs in detrusor and mucosa. (F) Venn diagrams of comparing upregulated and down regulated DEGs in detrusor and mucosa.

We first sought to test the impact of SCI on the bladder transcriptome and the extent to which differentially expressed genes (DEGs) separate different samples and treatment groups from each other. The heatmap and hierarchical clustering based on the top most variable genes in the whole dataset (18 samples) indicated two main clusters comprising detrusor and mucosa samples. All subgroups of SCI +/- inosine and controls were also clustered together indicating the impact of both injury and inosine treatment on the bladder transcriptome (Fig. 1C). The separation of detrusor from mucosa based on DEGs was also confirmed by principal component analysis (PCA) (Fig. 1D). In the PCA, detrusor samples from vehicle-treated SCI rats clustered further from detrusor samples from inosine-treated SCI rats, which clustered close to controls. Using the top 200 genes did not separate mucosa samples in different treatment groups when compared with detrusor. However, when detrusor samples were excluded from analysis, we observed a complete separation of control mucosa, vehicle-treated SCI mucosa and inosine-treated SCI mucosa (Fig. 1D, upper right panel) using the same 200-gene set.

In detrusor, SCI was associated with 1582 DEGs whereas in mucosa 564 DEGs were detected when compared to their respective controls (Fig. 1E&F). Of these DEGs, 1051/1582 were upregulated and 531/1582 were downregulated in detrusor of vehicle-treated SCI animals whereas 196/564 were upregulated and 368/564 were downregulated in mucosa of vehicle-treated SCI animals (Fig.1E, Suppl. Fig. 2A&B, 3A&B). In inosine-treated SCI animals, we identified 1740 DEGs in detrusor (1213 upregulated, 527 downregulated) and 287 DEGs in mucosa (181 upregulated, 106 downregulated) (Fig. 1E&F). Our analysis revealed that the expression level of 510 genes in detrusor and 172 genes in mucosa that were upregulated with SCI, was preserved after inosine treatment, showing expression comparable to that observed in control samples (Fig. 1E&F). Similarly, the expression level of 304 genes in detrusor and 322 genes in mucosa that were downregulated with SCI, was preserved with inosine treatment, with a level similar to that in control samples (Fig. 1E&F). Collectively, these genes are considered to be inosine-responsive.

Among the inosine-responsive genes, 29 were present in both detrusor and mucosa: 18 that were upregulated with injury, but maintained at control levels with inosine (*Rbm47, Rpl, Zfand4, Tmco4, Mid1ip1, Aspg AAABR07072078.1, Sh3yl1, Slc35d3, Slc9a4, Rcc, Slc15a2, Slc24a2, Tor3a, Coq7, Gltp, Sowaha, LOC688778*) (Fig. 1FG, Suppl. Fig. 4A &B, Suppl. Table 1-6) and 11 that were downregulated with injury, but maintained at control levels with inosine (*Cenpf, Galnt17, Lig4, Fam111a, Adamts12, Adam19, Ccdc88a, Aff2, Nusap1, Bco2, Rassf4*)(Fig. 1FG, Suppl. Fig. 4A & B, Suppl. Table 1-6). Among these genes, *Cenpf, Coq7, Rbm47, Rpl* and solute carrier family (*Slc*) genes were previously shown to be inosine-responsive in neurons [20, 21].

### Gene Ontology Analysis

Next we performed comparative over-representation analysis (ORA) using the Gene Ontology (GO) resource [22], to identify molecular functions (MFs), cellular components (CCs) and biological processes (BPs) differentially regulated in the bladder by SCI, and their sensitivity to inosine treatment. To determine the biological relevance of the molecular and cellular changes following SCI in the absence and presence of inosine treatment, we performed clustering of enriched terms and partitioned them into biologically-relevant groups. In the vehicle-treated detrusor, hierarchical clustering analysis of enriched molecular functions (MFs) revealed the top 3 clusters to be related to calcium signalling, syntaxin-1 and calmodulin binding, and extracellular matrix interactions (Suppl. Fig. 5A). Syntaxin-1 mediates the calcium-dependent trafficking of synaptic vesicles and subsequent neurotransmitter release at synapses [23, 24]. Following inosine treatment, syntaxin-1 binding and calcium binding-related functions were among the 37 SCI-induced MFs that were maintained at the control level, and therefore no longer enriched (Suppl. Fig. 5A & B), suggesting these are inosine-sensitive functions. These observations are consistent with the known neuroregulatory activities of inosine [14].

The hierarchical clustering analysis of regulated cellular components (CCs) in the detrusor revealed clusters associated with protein synthesis, structural integrity, cell-cell adhesion, and lysosomal compartments (Suppl. Fig. 6A). Inosine treatment resulted in changes in CC clusters, including upregulation of actin cytoskeleton, persistence of lysosomes, and alterations in cellular protrusions and adhesion structures (Suppl. Fig. 6B). These changes suggest potential remodelling and restoration of cellular organization and function in the detrusor. The hierarchical clustering analysis of regulated biological processes (BPs) in the detrusor identified clusters related to protein synthesis, cellular signalling, immune response, cell adhesion, and smooth muscle cell proliferation (Figure 2A). However, following inosine treatment, distinct changes were observed in the clustering patterns including DNA repair, and positive regulation of DNA-binding transcription factor activity. These findings suggest that inosine treatment may have elicited its effect partially by influencing DNA repair mechanisms and transcriptional regulation processes, while also impacting ion homeostasis and cellular communication (Figure 2B & C).

**Figure 2.**
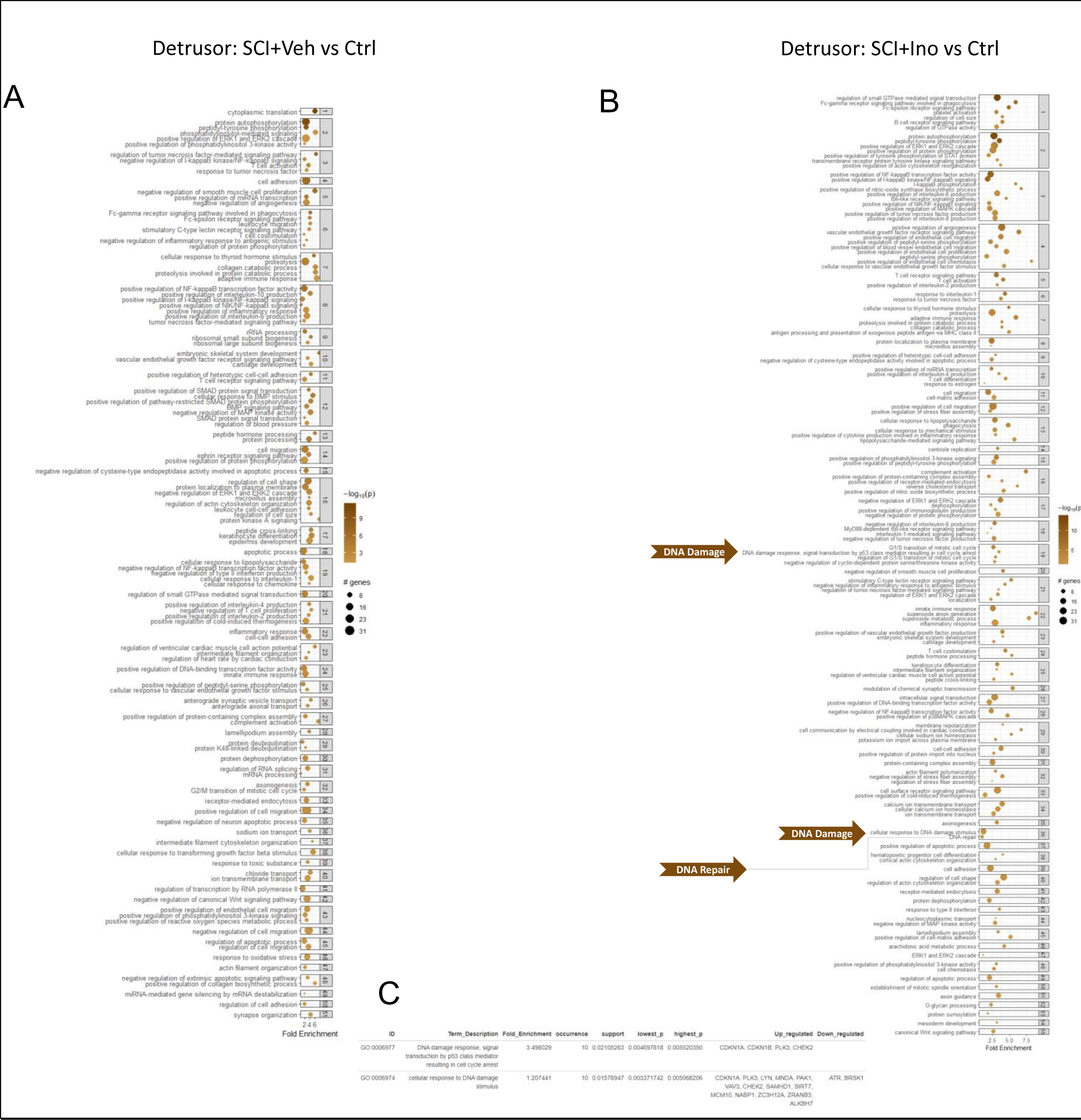
Hierarchically Clustered GO Biological Processes (BPs) Based on Regulated RNAs in Detrusor of Spinal Cord-Injured (SC) Rats. (A) Hierarchically Clustered GO Biological Processes (BPs) based on regulated RNAs in the detrusor of SCI rats treated with vehicle compared to controls. (B) Hierarchically Clustered GO Biological Processes (BPs) based on regulated RNAs in the detrusor of SCI rats treated with inosine compared to controls. The intensity or shade of brown in the bubbles represents the significance or enrichment of the Gene Ontology (GO) Biological Processes (BPs) term. Darker, more intense brown indicates higher significance or stronger enrichment of the term. The size of the bubbles corresponds to the number of genes associated with the particular GO BP term. Larger bubbles indicate that a greater number of genes in your dataset are associated with that specific BP term. (C) detailed table of enriched genes in the BPs related to DNA damage.

In the mucosa, analysis of MFs revealed clusters associated with protein kinase activity, protein degradation, cell cycle regulation, TGF-β signalling, post-transcriptional gene regulation, ion channel activity, and protein kinase signalling pathways following SCI, many of which were modulated with inosine treatment (Suppl. Fig. 7A & B). Inosine treatment showed potential modulation of these functions, indicating its involvement in cell cycle regulation, protein degradation, TGF-β signalling, gene expression control, ion channel activity, and protein kinase signalling. The analysis of CCs in the mucosa identified clusters representing diverse cellular structures, including extracellular matrix components, membrane structures, cytoskeletal elements, and muscle contractile units (Suppl. Fig. 8A & B). Inosine treatment influenced cellular component clusters associated with cell cycle regulation, gene expression control, neuronal morphology, cytoskeletal organization, intercellular communication, muscle contraction, and tissue integrity. Analysis of BPs in the mucosa revealed clusters associated with DNA damage response, cell cycle regulation, signalling pathways, and tissue repair processes (Figure. 3A & B). Inosine treatment significantly expanded the number of BPs and clusters, suggesting its impact on cell cycle regulation, cellular senescence, synaptic potentiation, receptor-mediated endocytosis, and regulation of membrane repolarization (Figure. 3B). Inosine treatment also influenced DNA damage response, regulation of DNA damage signalling, and cellular responses to DNA damage stimulus (Figure. 3C). These findings collectively demonstrate the diverse molecular and cellular changes induced by spinal cord injury and the potential therapeutic effects of inosine in the detrusor and mucosa. Inosine treatment has the potential to modulate calcium signalling, protein phosphorylation, transcriptional regulation, cellular adhesion, cell cycle regulation, DNA damage response, and other cellular processes.

**Figure 3.**
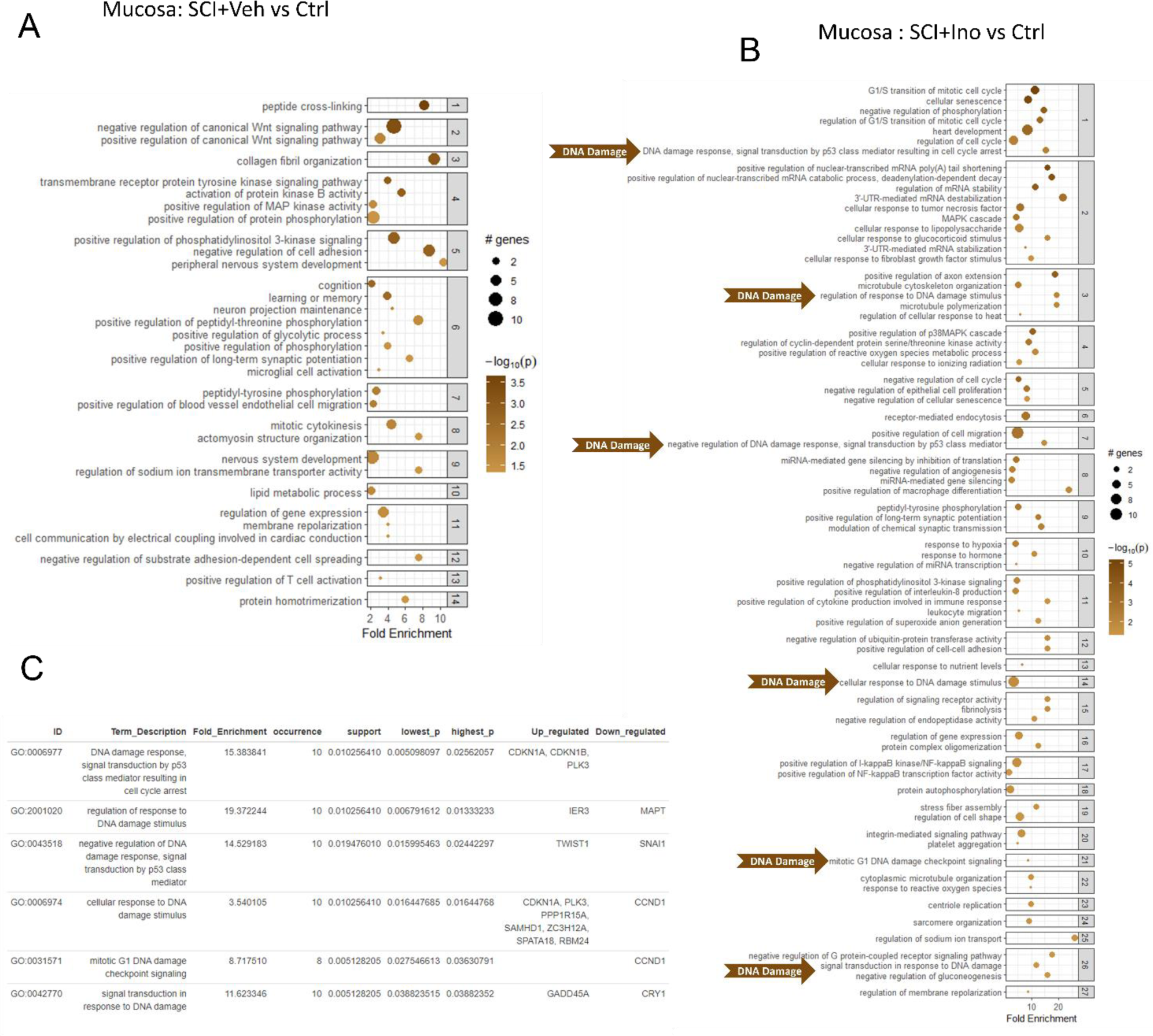
Hierarchically Clustered GO Biological Processes (BPs) Based on Regulated RNAs in Mucosa of SCI Rats. (A) Hierarchically Clustered GO Biological Processes (BPs) based on regulated RNAs in the mucosa of SCI rats treated with vehicle compared to controls. (B) Hierarchically Clustered GO Biological Processes (BPs) based on regulated RNAs in the mucosa of SCI rats treated with inosine compared to controls. (C) detailed table of enriched genes in the BPs related to DNA damage.

### Canonical Pathways Regulated by SCI

In order to identify regulatory pathways altered in the bladders of SCI rats, we next performed canonical pathway analysis in Ingenuity Pathway Analysis (IPA). The analysis combined ORA using GO and IPA (Fig. 2, 3 & 4). The top pathways in the detrusor of vehicle-treated SCI rats compared to controls included EIF2 signalling, Complement system, Axonal Guidance signalling, Synaptogenesis signalling, and mTOR signalling (Fig. 4A, Suppl. Fig. 9A). Consistent with its role in regulation of protein synthesis and translational control, the EIF2 signalling pathway was enriched for multiple genes encoding ribosomal proteins (Suppl. Fig. 15A), whereas the Complement system pathway harboured genes encoding complement factors and regulators (Fig. 4C). The emergence of Axonal Guidance and Synaptogenesis signalling pathways is consistent with the neuroplasticity evident in the bladder following SCI (reviewed in [25]). To further explore the regulatory landscape, we conducted an analysis to identify the genes most frequently represented within the top 20 regulated pathways in the detrusor of vehicle-treated SCI versus controls. Genes enriched in these pathways included *Rap2b, Prkd1, Adcy3, Adcy7, Camk4, Arpc1b and Itpr3*, suggesting their involvement in the molecular mechanisms underlying the response of the bladder to SCI (Fig. 4B&D, Suppl. Fig. 10, Suppl. Fig. 14A).

**Figure 4.**
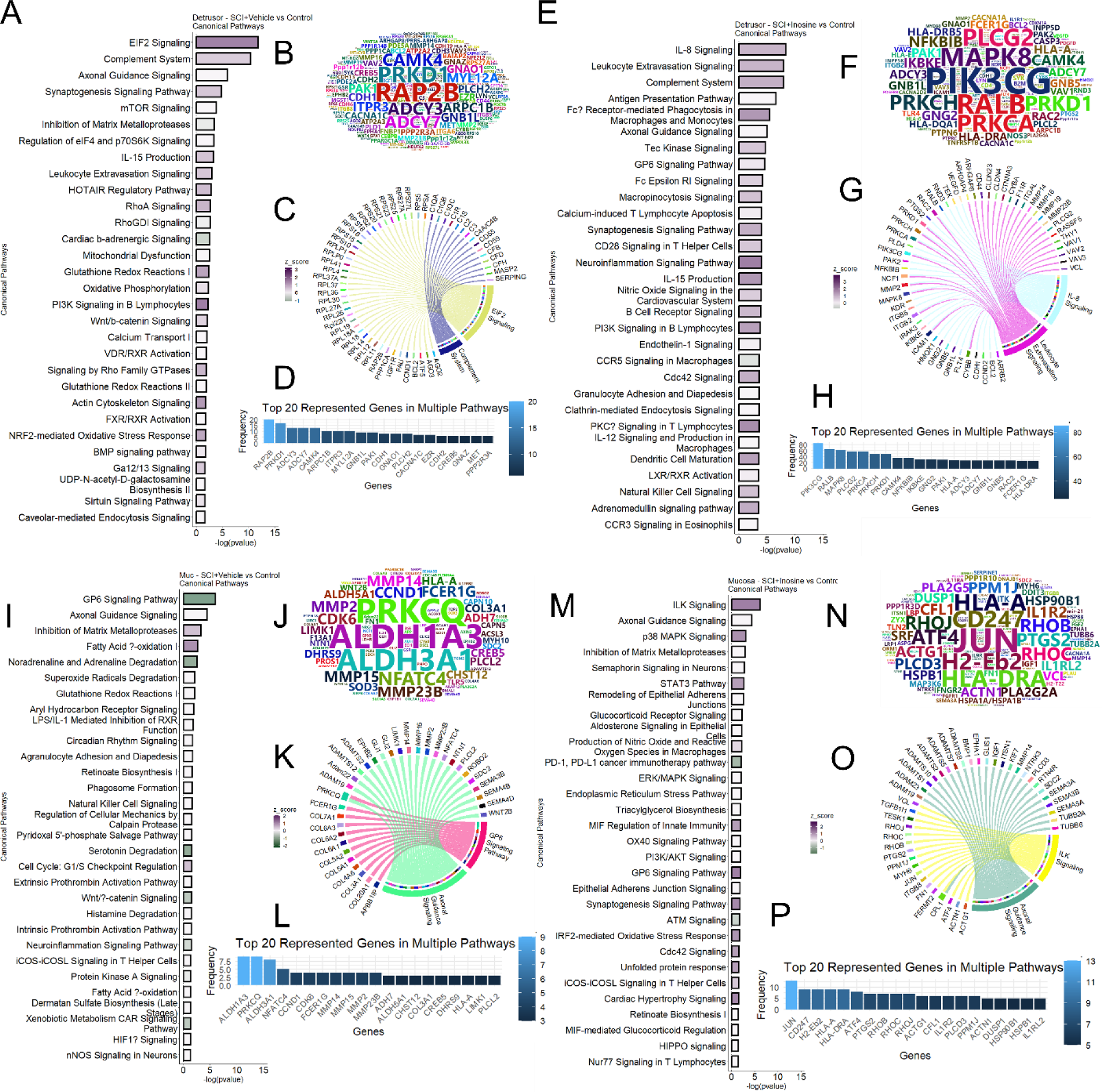
Regulated canonical pathways inferred by IPA. (A) Regulated pathways in detrusor following SCI treated with vehicle compared with control. Barchart of top 30 regulated pathways (B) Word cloud of most frequent genes regulated and enriched in the regulated pathway. (C) Circos plot of top 2 pathways (based on pvalue) and enriched genes. (D) Barchart visualizing the frequency of enrichment of top 20 enriched genes in pathways. (E) Regulated pathways in detrusor post SCI treated with inosine compared with control. Barchart of top 30 regulated pathways (F) Word cloud of most frequent genes regulated and enriched in the regulated pathways. (G) Circos plot of top 2 pathways (based on pvalue) and enriched genes. (H) Barchart visualizing the frequency of enrichment of top 20 enriched genes in pathways.(I) Regulated pathways in mucosa following SCI treated with vehicle compared with control. Barchart of top 30 regulated pathways (J) Word cloud of most frequent genes regulated and enriched in the regulated pathway. (K) Circos plot of top 2 pathways (based on pvalue) and enriched genes. (L) Barchart visualizing the frequency of enrichment of top 20 enriched genes in pathways.(M) Regulated pathways in mucosa post SCI treated with inosine compared with control. Barchart of top 30 regulated pathways (N) Word cloud of most frequent genes regulated and enriched in the regulated pathways. (O) Circos plot of top 2 pathways (based on pvalue) and enriched genes. (P) Barchart visualizing the frequency of enrichment of top 20 enriched genes in pathways.

Top pathways in the detrusor of inosine-treated SCI rats compared to controls included several related to inflammation and immune cell biology such as IL-8 Signalling, Leukocyte Extravasation Signalling, Complement System, Antigen Presentation pathway and Fcγ receptor-mediated phagocytosis in macrophages and monocytes (Fig. 4E & G). In agreement with these findings, inosine has been implicated previously in regulation of inflammatory cell signalling [26, 27]. The genes most frequently represented in top regulated pathways included *Pik3cg*, *Ralb*, *Mapk8*, *Plcg2*, *Prkca*, *Prkch*, *Prkd1*, *Camk4*, *Nfkbib* and *Ikbke* indicating their involvement in the regulation of identified pathways (Fig. 4F & H, Suppl. Fig. 11, Suppl. Fig. 14B). Notably, analysis of the pathways no longer enriched in detrusor following inosine treatment identified EIF2 signalling, RhoA and Signalling by Rho family GTPases, β-adrenergic signalling, Glutathione Redox Reactions and NRF2-mediated oxidative stress response (Suppl. Fig. 9A). Therefore processes regulated by such pathways following SCI, including protein synthesis, cytoskeletal regulation, and oxidative stress, appear to be sensitive to treatment with inosine.

The top regulated canonical pathways identified in the mucosa of vehicle-treated SCI rats compared to controls were GP6 signalling pathway, Axonal Guidance signalling, Inhibition of Matrix Metalloproteases, Fatty Acid β-oxidation, and Noradrenaline and Adrenaline degradation (Fig. 4I). Consistent with its role in wound healing, the GP6 Signalling pathway was enriched in multiple collagen-encoding genes as well as several inflammation-related genes, whereas the Axonal Guidance signalling pathway was enriched in multiple genes encoding regulators of extracellular matrix turnover such as *Adam* and *Mmp* family members (Fig. 4K). The genes most frequently enriched among the top 20 regulated pathways in mucosa of vehicle-treated SCI rats versus controls included *Aldh1a3, Prkcq, Aldh3a1, Nfatc4, Ccnd1, Cdk6, Fcer1g, Mmp14, Mmp15, Mmp2,* and *Mmp23b*. These genes signify their potential significance in the biological processes underlying the observed pathway alterations, highlighting their potential roles as key players in the mucosa in response to SCI (Fig. 4J&L, Suppl. Fig. 12, Suppl. Fig. 14C).

Pathway analysis of data from mucosa of inosine-treated SCI rats compared to controls identified ILK Signalling, Axonal Guidance Signalling, p38 MAPK Signalling, Inhibition of Matrix Metalloproteases, and Semaphorin Signalling in Neurons (Fig. 4M) as the most significantly enriched. Genes enriched in ILK Signalling included those encoding regulators of the cytoskeleton and contractility (*Actg1, Actn1, Cfl1, Fn1, Itgb8, Myh6, Rhob, Rhoc, Rhoj, Vcl)* as well as transcription factors (*Atf4, Jun*). The genes most frequently represented within the top 20 regulated pathways in the mucosa of inosine-treated SCI rats compared to controls included *Jun, Cd247, H2-Eb2, Hla-A, Hla-Dra, Atf4, Ptgs2, Rhob, Rhoc, and Rhoj* (Fig. 4N&P, Suppl. Fig. 13, Suppl. Fig. 14D). The top pathways no longer enriched in mucosa following inosine treatment included Inhibition of Matrix Metalloproteases, Fatty Acid Oxidation I, Noradrenaline and Adrenaline Degradation, Serotonin Degradation and Cell Cycle: G1/S Checkpoint Regulation (Suppl. Fig. 9B). While the GP6 Signalling Pathway still showed some level of enrichment, this was substantially reduced in mucosa following inosine treatment. These findings suggest that processes such as ECM turnover and neurotransmitter regulation in the mucosa are sensitive to inosine treatment following SCI.

### Identification of Master Regulators in Bladders of Spinal Cord Injured Rats

Following the identification of differentially regulated genes in detrusor and mucosa associated with SCI, we next aimed to identify master regulators that were predicted to drive the changes in gene expression observed with SCI, and to identify potential modulators of these. To accomplish these goals, we used causal approaches in Ingenuity Pathway Analysis [28] to predict the most significant upstream regulators and expose causal relationships associated with genes including regulators that are directly and indirectly connected to targets in our dataset.

Upstream Regulator Analysis (URA) of gene expression data from vehicle-treated SCI detrusor versus control predicted TP53 and MYC as the transcription regulators with the most significant p-values of overlap and activation Z-scores >2. Causal Network Analysis (CNA), which enabled us to gain more extensive insights into putative master regulators, predicted TRRAP as the most significant transcription regulator (Z-score, 2.838; p-value of overlap, 1.39E-34) from DEGs from vehicle-treated SCI detrusor versus control. TRRAP encodes a member of the PIKK family of PI3K-related kinases which includes DNA-PKcs, ATM and ATR, that senses and responds to DNA damage [29]. TRRAP is known to interact with TIP60, which was predicted as significantly activated in our data (Z-score, 3.619; p-value of overlap, 6.3E-30) and has been implicated in DNA repair [30].

We also used CNA to identify potential modulators of the observed gene expression changes. Since our goal was to identify agents that could form the basis of treatments, we focused on endogenous chemicals, with the rationale that these would tend to be well tolerated in vivo (Fig. 5A-D). Analysis of vehicle-treated SCI detrusor versus control predicted inosine as the most significant endogenous chemical (Z-score, −1.53, p-value of overlap, 1.14E-29)(Fig. 5A). The negative z-score indicated that inosine was predicted to inhibit the corresponding networks. By interrogating the putative targets we inferred that inosine acts upstream of poly-ADP ribose phosphorylase-1 (PARP1) within the detrusor following SCI. The molecules downstream of PARP1 were found to be activated, leading us to conclude that PARP1 is activated in the bladder following SCI (Fig. 5C). PARP1 was predicted to activate *Ar, Atf2, Akt, Creb1, Elk1, Erk, Gata3, H2ax, Ikbkg, Jnk, Mapk8, Mmp9, Mybl2, Nfkb (Complex), Nos2, P38 Mapk, Prkdc, Rela, Rar, Stat3, and Tp53.* Additionally, PARP1 was predicted to be involved in regulation of several molecules, notably *Atf2, Akt, Creb1, Elk1, Erk, H2ax, Mybl2, Rela, Stat3, and Tp53* (Fig. 5C). Interestingly, CNA also predicted inosine to activate a distinct downstream network (Z-score, 4.264; p-value of overlap, 5.37E-16), that was non-overlapping with the PARP1-regulated network (Fig. 5D). The predicted targets in this inosine-activated network include complement factors, the annexins *Anxa1* and *Anxa3*, and genes encoding actin-binding proteins such as Arpc1b and Lcp1. The prediction of both inosine-inhibited and inosine-activated networks by CNA suggests that SCI itself modulates inosine metabolism in the bladder.

**Figure 5.**
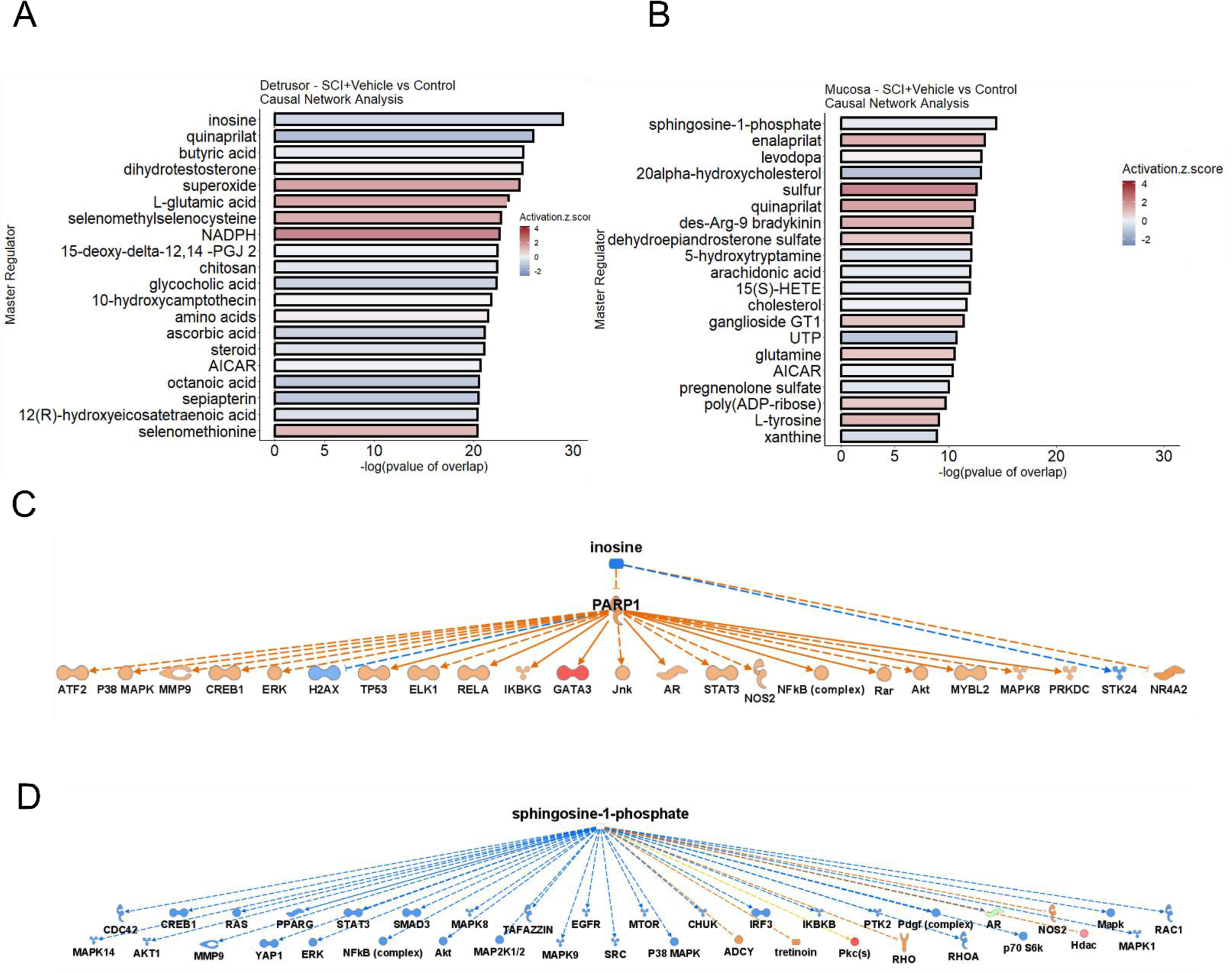
Causal Network Analysis. (A) Causal network analysis in detrusor of SCI rats treated with vehicle compared to uninjured controls. (B) Causal network analysis in mucosa of SCI rats treated with vehicle compared to uninjured controls containing top 25 regulator molecules (including inosine and PARP1). (C) Causal network of top endogenous chemical in the detrusor of SCI rats treated with vehicle. (D) Causal network of top endogenous chemical in the detrusor of SCI rats treated with vehicle containing top 39 regulator molecules.

URA of vehicle-treated SCI mucosa versus control predicted MYC, as well as SPDEF, SMAD7 and PPARGC1A as transcription regulators with the most significant p-values of overlap and activation Z-scores >2. CNA of DEGs from vehicle-treated SCI mucosa versus control predicted one of the most significant regulators in mucosa to be the DDB1/CUL4/RBX1 complex, which is known to regulate DNA damage response signalling [31]. Among the endogenous chemicals, sphingosine-1-phosphate (S1P) emerged as the most significantly altered (Z-score, −0.659; p-value of overlap, 3.97E-15)(Fig. 5B). sphingosine kinase 1, which generates S1P from sphingosine, was also predicted to be inhibited (Z-score, −2.157; p-value of overlap, 1.9E-15) in agreement with the predicted decrease in S1P. Consistent with findings in detrusor, S1P has been linked in prior studies to modulation of or response to DNA damage in neurons [32]. Bioinformatics analysis to predict functional relationships between S1P and downstream genes revealed that S1P inhibits *Adcy, Hdac,* and *Mtor.* In addition, S1P was predicted to be involved in the phosphorylation of multiple molecules including *Akt1, Chuk, Creb1, Egfr* and *Erk* (Fig. 5D).

### Validation of bioinformatics analysis

To validate findings from URA and CNA, and test the prediction that DNA damage may act as a driver of the observed gene expression changes in the bladder wall following SCI via PARP-1, we stained bladder tissues from rats without and with SCI treated with vehicle or inosine for gamma-H2AX (γH2AX), a marker of DNA damage and inducer of PARP-1 activation, and poly-ADP ribose (PAR), the product of PARP-1 activation. We found that both γH2AX (Fig. 6A) and PAR (Fig.6B) were induced by SCI compared to control, in both detrusor and mucosa. Consistent with the prediction that inosine inhibits PARP-1, staining for γH2AX and PAR was attenuated in tissues from SCI rats treated with inosine. Additionally, we verified the presence of PAR staining in tissues obtained from patients with neurogenic bladder (Fig. 6C).

**Figure 6.**
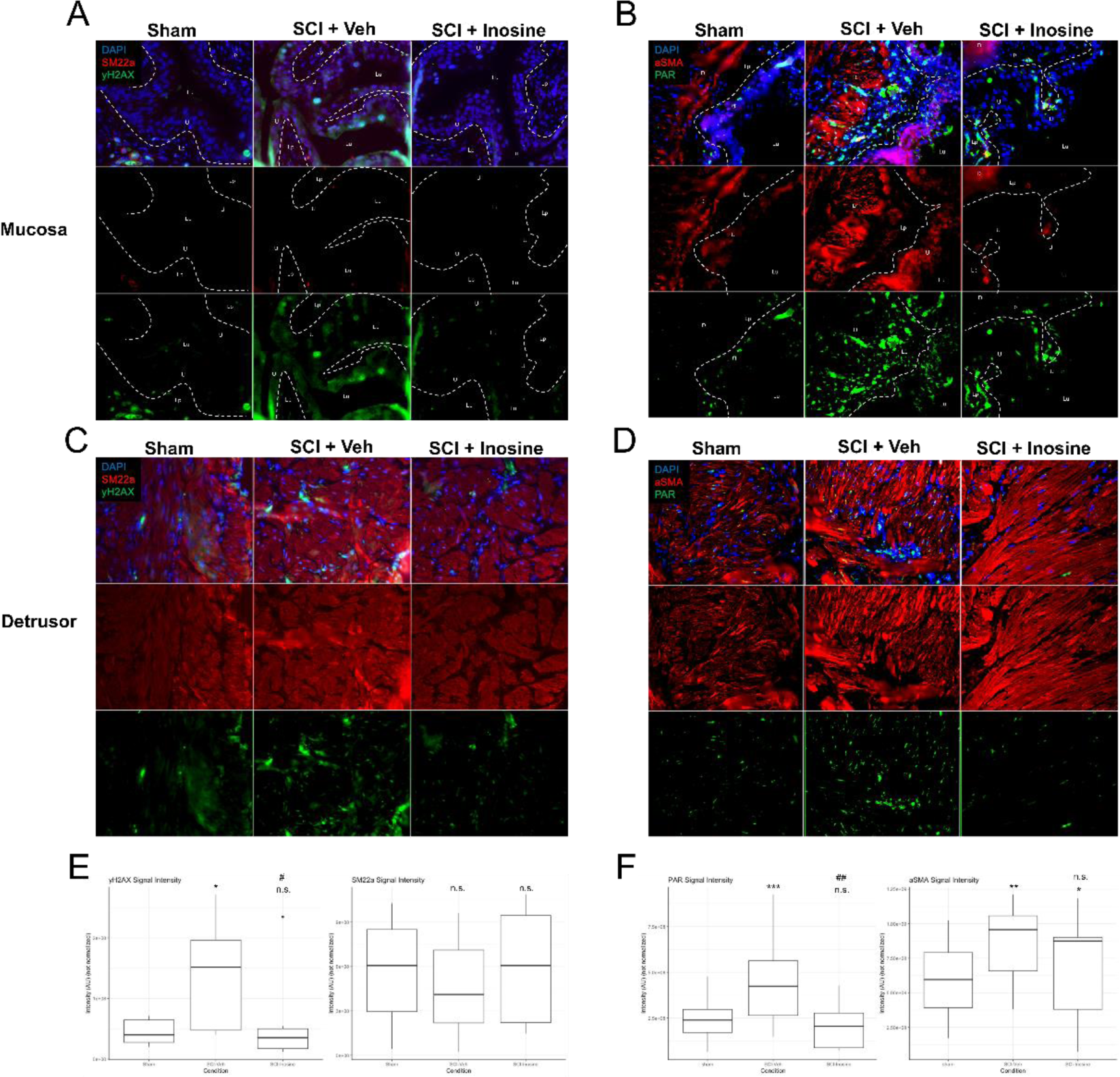
Immunostaining for CNA and pathway validation. (A) Bladder sections from SCI rats were stained with primary antibodies to γH2AX (green) and SM22α (red) (A, C) or PAR (green) and αSMA (red) (B, D). Data are representative of 4 fields captured from each of 3 biological replicates. Note the increase in γH2AX (red) and PAR (green) signal with injury (SCI-Veh vs control) and the significant reduction in γH2AX (red) or PAR (green) signal following inosine treatment compared to vehicle treated SCI (SCI-Inosine vs SCI-Veh) (E, F). Dotted lines separate the lamina propria (Lp) and the urothelium (U). Lu indicates the lumen and the D indicates the detrusor. ANOVA and Tukey Honest Significant Differences were used to calculate pvalues. * p 0.05, ** p 0.01, comparing to the Sham. # p 0.05, ## p 0.01, comparing to the SCI.

### Proteomics analysis

In addition to transcriptomics analysis, we also performed quantitative proteomics analysis on full thickness bladder samples obtained from rats at 8 weeks following SCI compared to uninjured controls. Analysis of differentially expressed proteins using IPA revealed 296 canonical pathways that were significantly regulated in response to SCI. Notably, enrichment of Regulation of eIF4 and p70S6K Signalling, Mitochondrial Dysfunction and EIF2 Signalling suggest alterations in protein synthesis and energy production, potentially reflecting cellular responses to injury. Additionally, identification of NRF2-mediated Oxidative Stress Response and Oxidative Phosphorylation as enriched pathways point to heightened oxidative stress and potential mitochondrial dysfunction, whereas identification of Sirtuin Signalling Path ay and PPARα/RXRα Activation suggest alterations in inflammation and metabolism following SCI (Fig. 7A). Consistent with the impact of SCI on the nervous system, we also identified a number of pathways related to neuronal function. These include the Reelin Signalling in Neurons pathway, which is closely tied to the regulation of neurogenesis, neuronal migration and synaptic plasticity [33] and the Axonal Guidance Signalling pathway implicated in regulating neuronal guidance during development, as well as CDK5 Signalling which plays a pivotal role in orchestrating neuronal cell cycle progression, synaptic plasticity, and neuronal development [34]. CDK5 has been implicated in the response to DNA damage [35] consistent with our demonstration of increased γH2AX staining in the bladder following SCI. We also observed significant enrichment of MAPK1, MAPK3, MAP2K1, and MAP2K2 proteins within the regulated pathways (Fig. 7B & C), consistent with the identification of the ERK/MAPK Signaling pathway among the top regulated canonical pathways and indicating the potential activation of MAPK signalling in response to SCI. In line with this level of enrichment, MAPKs were highly involved in a majority of the top 20 regulated pathways including Regulation of eIF4 and p70S6K Signalling, NRF2-mediated Oxidative Stress Response, Insulin Secretion Signalling Pathway, Reelin Signalling in Neurons, Signalling by Rho Family GTPases, and mTOR Signalling (Fig. 7D).

**Figure 7.**
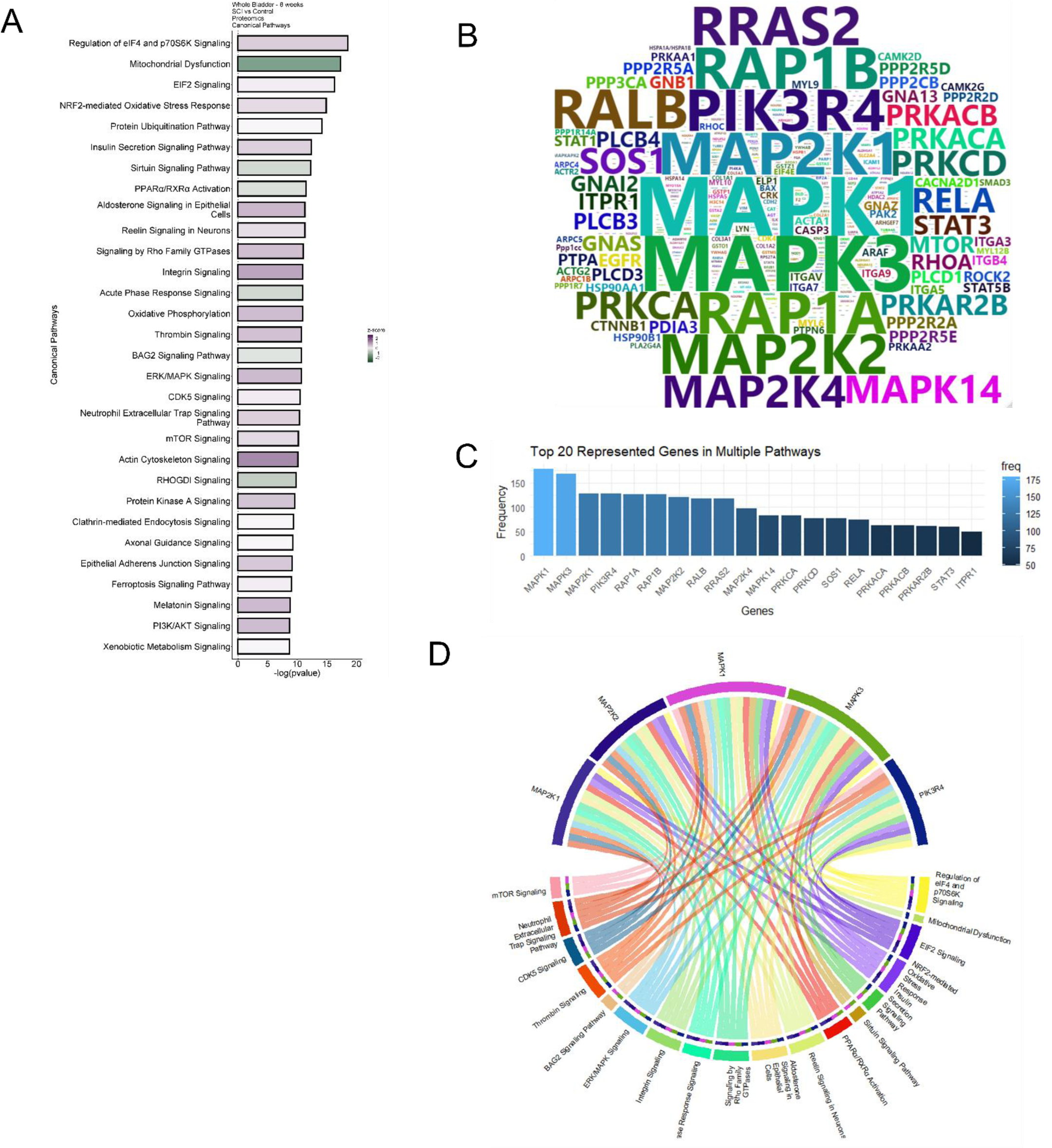
Regulated canonical pathways inferred by IPA in 8 weeks proteomics. (A) Barchart of top 30 regulated pathways in whole bladder of 8 weeks post SCI compared with control. (B) Word cloud of most frequent proteins regulated and enriched in the regulated pathways. (C) Barchart showing the frequency of top 20 most recurrent protein enriched in pathways. (D) Circos plot of top 5 most recurrent enriched proteins in the regulated pathways and the corresponding regulated pathways.

### Integration of multi-omics data

In order to gain a greater understanding of pathway regulation in the context of SCI, we compared the significantly regulated pathways identified from proteomics analysis of full thickness bladder samples with those identified from transcriptomics analysis of separated detrusor, and mucosa from bladders at 8 weeks following SCI, and identified considerable overlap between them (Suppl. Fig. 16). Ten pathways were common among all three datasets, including Axonal Guidance Signalling, GP6 Signalling Path ay, Gl tathione Redox Reactions and nt/β-catenin Signalling. Furthermore, 15 pathways were matched exclusively between those identified by proteomics of whole bladder and those enriched in the mucosa, as determined by RNA sequencing. These included HIF1α Signalling, Superoxide Radicals Degradation, Fatty Acid β-oxidation and pathways related to inflammation and immune cell function. In the detrusor, 49 pathways exhibited exclusive intersection with those identified by proteomics of the whole bladder and those identified by RNA sequencing of detrusor. Taken together, this integrated analysis identified important regulators of the bladder tissue response to SCI as well as potential therapeutic targets, but also uncovered distinct responses within the detrusor and mucosa. These latter differences, together with the differing sensitivity of detrusor and mucosa to inosine, warrant further investigation, as they are likely to reveal cell type-selective functions in the context of injury.

## Discussion

Previous findings from us and others have demonstrated beneficial activity of inosine treatment on bladder function in the setting of SCI in preclinical models [13, 16, 36]. In this study, we described the molecular landscape of the bladder following SCI in rats and applied a multi-omics approach to obtain novel insights into both neurogenic bladder pathobiology and mechanisms of inosine action. This study provides the first demonstration of PARP activation in the bladder wall in response to SCI, and its attenuation with inosine treatment.

Among the molecular alterations observed in our model, the emergence of DNA damage following SCI was particularly striking. Pathways linked to oxidative stress, a known inducer of DNA damage, were also enriched with SCI. These included NRF2-mediated oxidative stress response, mitochondrial dysfunction, glutathione redox reactions and HIF-1α signaling. Oxidative stress is known to arise early in the bladder after SCI. Loss of normal innervation to both the bladder and external urethral sphincter results in detrusor-sphincter dyssynergia, with the ensuing functional obstruction leading to increased pressure within the bladder during filling, thereby altering blood flow and resulting in transient ischemia/hypoxia and oxidative stress [37, 38]. Published reports have described increases in reactive oxygen species (ROS) in urothelial cells as early 3 days after injury [39]. In support of a causative relationship between oxidative stress and DNA damage following SCI, we also observe co-occurrence of ROS and γH2AX signals in the same cells as early as 3 days following SCI in mice (AHG, unpublished observations), with DNA damage persisting for at least 4 weeks. Reports in the literature have shown that persistent DNA damage can in turn evoke oxidative stress [40] suggesting the existence of a feed-forward loop that drives pathologic changes in the bladder in the setting of ongoing obstruction.

Of particular interest to us was the prevention of DNA damage in tissues of inosine-treated animals. Inosine is a known anti-oxidant related in part to its metabolism to urate, a scavenger of reactive oxygen and reactive nitrogen species (reviewed in [14]). In agreement with these observations, one of the genes identified as inosine-sensitive in both detrusor and mucosa was *Coq7* (coenzyme Q7), which encodes the mitochondrial enzyme CLK-1, implicated in cellular respiration. *Coq7* was induced in the bladder in response to SCI but maintained at control levels in tissues from inosine-treated animals. In previous studies, decreased CLK-1 levels were associated with attenuation of mitochondrial oxidative stress in models of accelerated aging [41]. Moreover, the neurodegeneration-inhibiting agent clioquinol was found to act through inhibition of CLK-1 in cell and animal models [42]. CLK-1-mediated signaling has also been shown to regulate stress responses in mitochondria, including the metabolism of reactive oxygen species and the unfolded protein response [43], both of which have been implicated in bladder dysfunction of both neurogenic and non-neurogenic origin [39, 44, 45].

Consistent with the enrichment of oxidative stress-associated pathways and DNA damage, SCI was also associated with activation of PARP-1, which itself was inhibited with inosine treatment. Purines, including inosine, have been shown previously to inhibit PARP-1 activity under cell free conditions and in cultured macrophages exposed to peroxynitrite [46] and PARP inhibition has led to attenuation of oxidative injury in other organs [47]. Interestingly, Virag and Szabo showed that the inosine metabolite, hypoxanthine, was more potent at inhibiting PARP than inosine and also displayed greater cytoprotection of macrophages exposed to oxidative stress compared to inosine [46]. These observations are in contrast to findings in the lower urinary tract where hypoxanthine has been associated with damage to the bladder with age [48, 49]. Consistent with a causative role for hypoxanthine, inhibition of purine nucleoside phosphorylase (PNPase), the enzyme that converts inosine to hypoxanthine, in aged rats, was associated with improvement in multiple aspects of bladder morphology and function [48]. These included mitochondrial damage, oxidative stress and voiding dysfunction, all of which are also evident in neurogenic bladder following acute SCI in spite of the distinct etiology. Analysis of our proteomics data revealed a significant increase in levels of PNPase in bladder tissue from rats with SCI (Suppl. Fig. 16 B), consistent with the possibility of increased inosine metabolism to hypoxanthine and its associated deleterious impact on tissue as a consequence of injury. Taken together, these findings support the concept that bladder homeostasis is influenced by the balance between protective and toxic purines as exemplified by inosine and hypoxanthine, respectively [48]. Furthermore they reinforce the idea that increasing inosine levels in the urinary tract, whether by exogenous administration as described by us [13] or by inhibition of inosine metabolism to hypoxanthine [48] represents an effective therapeutic approach to prevent the deleterious consequences of injury to the bladder.

A major strength of this study is the integrated analysis of transcriptomics and proteomics data which identified several pathways that were regulated in a concordant manner following SCI. These included EIF2 signaling, NRF2-mediated oxidative stress response, RhoA Signaling, Actin Cytoskeleton Signaling, GP6 Signaling, Protein Kinase A and Wnt/β-catenin Signaling, a number of which were attenuated in tissues from inosine-treated animals. Alterations in EIF2 signaling are central to the integrated stress response in cells and tissue exposed to injury, that mediates compensatory changes in protein synthesis thereby enabling recovery and repair. However, although beneficial early after injury, sustained activation of the ISR has been associated with deleterious consequences in a variety of disease settings, leading to a search for strategies to inhibit EIF2 signaling to prevent the ISR from becoming maladaptive (reviewed in [50]). EIF2 signaling has been implicated in pathological changes in the brain and spinal cord neurons following injury, with eIF2α phosphorylation, central to the ISR, increased early after injury in parallel with emergence of morphological and functional deficits [51, 52]. In contrast, inhibition of EIF2 signaling using the small molecule ISRIB was associated with both histological and functional improvements. Thus, inhibition of EIF2 signaling with inosine may be predicted to achieve similar benefits in neuroprotection in the bladder in agreement with our prior findings [13].

There are a number of limitations in our study. All of our analyses were performed in male rats to enable comparison to previous studies conducted by our group [13, 16, 17], potentially limiting applicability of the findings to both sexes. We also lack proteomics data from separated detrusor and mucosa, such that we cannot assess the concordance between transcriptome and proteome in these tissues. Nonetheless, we were encouraged by the substantial overlap in commonly regulated pathways predicted from transcriptomics and proteomics analysis, which suggested strong agreement between them.

### Conclusions

In this study we present a comprehensive systems biology analysis of the bladder following traumatic SCI in a preclinical rat model. Our integrated transcriptomic and proteomic analysis highlights inosine’s role in modulating key signaling pathways, including EIF2 signaling, NRF2-mediated oxidative stress response, and nt/β-catenin signaling, essential for cellular stress responses and recovery. We also unveil the systemic impact of inosine following neurogenic injury, that extends beyond attenuation of DNA damage and PARP activation to the preservation of key biological networks and pathways. This rebalancing of molecular pathways by inosine underscores the potential of systems biology approaches to investigate complex neurogenic disorders and suggests new therapeutic possibilities, such as the repurposing of FDA-approved PARP inhibitors, for managing trauma-induced neurogenic dysfunction. This study not only contributes to our understanding of the molecular landscape following traumatic injury but also opens avenues for targeted, effective treatments grounded in systems biology.

## Materials and Methods

### Ethical Approval

The animal experiments conducted in this study were performed in strict accordance of the recommendations provided in the Guide for the Care and Use of Laboratory Animals of the National Institutes of Health. The experiments were approved by the Animal Care and Use Committee at Boston Children’s Hospital (protocol # 16-08-3256R).

### Creation of spinal cord injury model in rats

Complete spinal cord transection at T8 was performed in male Sprague Dawley rats (6-7 weeks of age, ∼250 grams, Charles River Laboratories, Wilmington, MA) under isoflurane anesthesia essentially as described [17]. Using spinal bony and vascular landmarks, the spinal cord was exposed at T8 and sharply transected. Gelfoam (EthiconTM) was interposed between the proximal and distal spinal cord ends to promote haemostasis; the dura was not closed. Paraspinal muscles and skin were closed in 2 layers. Post-operative pain and infection prophylaxis were managed with meloxicam analgesia (5 g/ l, s.c, every 24 h for 3 d) and Baytril (100 g/ l at .5 g/ g, s.c.), respectively. D ring the period characterized by bladder areflexia, bladders were emptied every 12 h by manual expression until animals demonstrated the ability to void independently, typically within 10-14 d. Rats were monitored daily for signs of urine scald, phimosis, infection, or other systemic illness, and weighed. In some animals with SCI, inosine (225 mg/kg/d) or a corresponding volume of vehicle (12 mM sodium bicarbonate buffer, pH 9.2) was administered by intraperitoneal injection daily for 8 weeks. In a separate experiment, animals were maintained for 2, 8 or 16 weeks after SCI. At the end of the treatment period, animals were euthanized via CO_2_ inhalation and tissues harvested in ice-cold, oxygenated Kreb’s b ffer (120 mM NaCl; 5.9 mM KCl; 25 mM NaHCO_3_; 1.2 mM Na_2_H_2_PO_4_; 1.2 M MgCl • 6H_2_O; 2.5 mM CaCl_2_; 11.5 mM dextrose). Bladders were weighed and either microdissected into detrusor and mucosa, or full thickness specimens prior to flash freezing in liquid nitrogen and storage at −80 C, or fixed in neutral-buffered formalin and processed for embedding in paraffin.

### Immunofluorescence staining and analysis

Full-thickness tissue sections (5 µm) were cut and subjected to immunofluorescence staining as described [13]. Primary antibodies included: gamma-H2AX (Abcam, 1:100 dilution), PAR (CST, 1:100 dilution), α-SMA (MilliporeSigma, 1:100 dilution) and SM22α (CST, 1:100 dilution). Sections were incubated with species-specific secondary antibodies conjugated to Alexa-488 and Alexa-594, with nuclei counterstained with DAPI. Signals were visualized on an Axioplan-2 microscope (Zeiss) and representative images captured using Axiovision software. An average of 6 fields of view were captured for each of 3 biological replicates per condition. Images were quantified using an in-house developed macro that quantifies the integrated density of the signal for each channel. A one-way ANOVA with was performed followed by a multiple pairwise-comparison between the means of groups using the Tukey Honest Significant Differences. Adjusted p values were used to report the significance of the differences.

### RNA isolation and sequencing

Tissues were lysed in TRIzol (Thermofisher) using Fast Prep Lysing matrix D beads (MP Biomedical). RNA was isolated and purified from the tissue lysates using miRNeasy columns (Qiagen) according to the an fact rer’s instr ctions. Multimodal RNA quantification and quality control was performed using NanoDrop (Thermofisher), Qubit RNA Broad-Range Assay kit (Thermofisher), and Agilent 2100 Bioanalyzer (Agilent). The ribodepletion method of library preparation for RNA sequencing was performed using the Total-RNASeq RiboErase Kit (KAPA Biosystems). All RNA samples were loaded two separate 2-lane flow cells and sequenced using an Illumina HiSeq 2500 system generating 51 bp paired-end reads. Read counts were combined from the two flow cells to obtain total read counts for each sample. Read count data was converted from BCL to FASTQ filetype using Bcl2fastq conversion software (Illumina).

#### Differential Gene Expression

Differential gene expression analysis of count data was performed using the Bioconductor R package DESeq2. Variance of the data was stabilized with logarithmic transformation (log2) of the normalized counts. Genes were considered to be differentially expressed if adjusted p-value <0.05. Gene ontology (GO) term identification was performed using gprofiler (http://biit.cs.ut.ee/gprofiler/) and clusterprofiler (https://bioconductor.org/packages/ release/bioc/html/clusterProfiler.html). Gene pattern clustering was performed using the DEGreport tool.

### Scatterplots of logFC vs logCPM

The ggplot R package was used to create the scatterplots of logFC vs logCPM. The code generates a scatter plot of log fold change (logFC) versus log counts per million (logCPM) of the top 100 differentially expressed genes between two groups. The X-axis represents the logarithmic fold change and the Y-axis shows the logarithmic count per million (CPM), which is a measure of the expression level of the genes in each sample. Additionally, red vertical lines are drawn at logFC = −2 and logFC = 2, and a red horizontal line is drawn at logCPM = 5. In this plot, each point represents a gene, and its position is determined by its fold change and CPM values. The points can be divided into different quadrants based on their location relative to the X and Y axes. The genes in the upper right quadrant are the upregulated genes, with both high fold change and high expression levels, while those in the lower left quadrant are the downregulated genes, with both low fold change and low expression levels. The genes are colored based on their logFC values.

### Heatmap and clustering

Hierarchical clustering and heatmaps were created using the “heatmap2” function available in the R package “Gene-E”. To construct the hierarchical clustering, we first computed a pairwise correlation matrix between the items, employing the Pearson correlation method. This correlation matrix was then transformed into a distance matrix. Subsequently, we performed clustering based on this distance matrix using the average linkage method, which utilizes the average distance for distance matrix calculation. The heatmaps were generated to visualize the hierarchical clustering results, specifically focusing on the differentially expressed genes (DEGs) within various comparison groups. DEGs were defined as those with a log2 fold change greater than or equal to ±0.5, a P-value less than 0.05, and read counts exceeding 1 count per million (cpm).

### Gene Ontology analysis

The workflow to calculate the fold enrichment in GO Molecular Function (MF), Cellular Compartment (CC) and Biological Process (BP), included taking a data frame of 3 columns containing: differentially expressed gene symbols, log2FC and p values. We then mapped these genes onto a protein-protein interaction network, which served as a reference database. The outcome of this analysis yields specific terms that exhibit enrichment of our Differentially Expressed Genes (DEGs). These enriched terms are referred to as “active subnetworks” and are characterized by the following columns: **ID:** ID of the enriched term; **Term_Description:** Description of the enriched term; **Fold_Enrichment:** Fold enrichment value for the enriched term (calculated using ONLY the input genes); **Occurrence:** the number of iterations that the given term was found to enriched over all iterations; **Support:** the median support (proportion of active subnetworks leading to enrichment within an iteration) over all iterations; **Lowest_p:** the lowest adjusted-p value of the given term over all iterations; **Highest_p:** the highest adjusted-p value of the given term over all iterations; **Up_regulated:** the up-regulated genes in the input involved in the given term’s gene set; **Down_regulated:** the down-regulated genes in the input involved in the given term’s gene set.

### Clustering GO Terms

Clustering can be performed via multiple different methods such as centroid, hierarchical or fuzzy method using the pairwise Cohen’s kappa statistics (a chance-corrected measure of co-occurrence between two sets of categorized data) matrix between all enriched terms. Based on biological context and o tco e of those ethods, e chose to se hierarchical cl stering of the ter s (sing 1−κ as the distance metric) and determined the optimal number of clusters by maximizing the average silhouette width. After clustering, we plot the summary enrichment chart was plotted and the enriched terms were displayed in clusters (Fig. 2 &3, Fig. Suppl. 5-7). This analysis enabled us to identify functionally related clusters, as well as prioritize and focus on specific functional clusters that are most relevant to the research question.

### Quantitative Proteomics analysis

Tandem mass tag (TMT)-based quantitative proteomics analysis was conducted essentially as we previously described [53]. Proteins were extracted from tissue samples by homogenizing tissue in SDS-containing lysis buffer (80 mM Tris-HCl, 4% SDS, 100 mM DTT, pH7.4). Protein concentrations were determined by using a Pierce 660nm Protein Assay kit (Thermo Scientific). From each sa ple, 60 μg proteins were alkylated by 55 mM iodoacetamide and digested with trypsin at 1:100 enzyme-to-protein ratio using the filter-aided sample preparation method [54]. Tryptic peptides were labeled with 11-plex TMT reagents in parallel, merged into one sample, desalted using C18 spin columns (Thermo Scientific), and dried down in a SpeedVac (Thermo Scientific). Each set of TMT11plex-labeled peptide mixture was fractionated into 48 fractions by high-pH liquid chromatography and concatenated into 16 fractions by combining fractions 1, 17, 33; 2, 18, 34; and so on. The concatenated fractions were concentrated in a SpeedVac and stored at −80°C. LC-SPS-MS3 analysis was performed on an EASY-nLC 1200 connected to an Orbitrap Fusion Lumos mass spectrometer (Thermo Scientific). Each fraction of TMT11plex-labeled peptides was resuspended in 0.2% formic acid, and about 1 µg peptide was loaded onto a 2-cm trap column and separated by a 50-cm EASY-Spray column (Thermo Scientific) heated to 55°C, using a 3-h gradient at a flow rate of 250 nL/min. The parameter settings for FTMS1 include orbitrap resolution of 120K, scan range of m/z 350-1400, maximum injection time of 100 ms, AGC target of 5E5, RF lens of 30%, data type of centroid, chage state of 2-5, dynamic exclusion for 60 s using a mass tolerance of 7 ppm, and internal calibration using 371.10123. The parameters for ITMS2 include mass range of 400-1400 m/z, 10 dependent scans, isolation window of 0.4 m/z, CID collision energy of 35%, maximum injection time of 120 ms, AGC target of 2E4, and data type of centroid. The parameters for MS3 include scan range of 100-1000 m/z, maximum injection time of 150 ms, AGC target of 2.5E5, HCD collision energy of 55%, and data type of centroid.

The acquired RAW files were searched against the Uniprot_Rat database (released on 07/18/2018) comprising canonical and isoform protein sequences (47,943 entries) with MaxQuant (v1.6.0.16) [55]. Carbamidomethyl (C) was set as fixed modification, while oxidation (M), acetyl (protein N-term), and deamidation (NQ) were set as variable modifications. The mass tolerance was 20 ppm for first search peptide tolerance, 4.5 ppm for main search peptide tolerance, and 0.5 Da for MS/MS match tolerance. The quantification type was set as Reporter ion MS3, the isobaric labels were TMT11plex, and the reporter mass tolerance was 0.003 Da. A standard false discovery rate of 1% was used to filter peptide-spectrum matches as well as peptide and protein identifications [53–55].

Because the total number of samples (20) in our proteomics experiment exceeded the maximum number for one set of isobaric tag reagents (11), we ran two TMT 11-plex experiments. In this case, each TMT experiment contained a pool of all 20 samples. The channels pooled reference mixture were used to match the protein reporter ion intensities between TMT experiments. This was done by dividing impurity corrected TMT intensity of each ID by the intensity value of its TMT pool, resulting in a pool normalization factor and then this factor was multiplied to the impurity corrected TMT intensities for that ID (for all samples). Further, we reasoned that there might be many sources of variation in addition to the biological differences between groups due to capacity of TMT experiment for running all samples; therefore, we designed a normalization pipeline to minimize the errors imposed by running samples in two batches. For TMT labeling, since we used the same amount of digested protein labeled for each channel, the liberated reporter ion signals were the proxies for protein abundance, so the sum of the reporter ions in each channel was a proxy for the total amount of protein and the total signal per channel was checked for consistency. We have compared two normalization methods. First, we performed normalization to abundant proteins and compared it with the second method where we used internal reference scaling (IRS) methodology [56] since it has been reported to be capable of correcting the random MS2 sampling that occurs between TMT experiments.

### Circos plots

We utilized Circos plots for the visualization and analysis of top canonical pathways and their corresponding enriched genes or proteins, with the analysis conducted sing the “circlize” pac age in R. The selection of top canonical pathways was based on the negative logarithm of p-values (Neg.log.p_value>1.3). To prepare the data for visualization, we transformed it by splitting the molecules within selected pathways and creating new rows for individual molecules. To establish the connections between pathways and molecules, we constructed a links data frame, serving as a representation of these relationships. Setting up the chord diagram visualization involved configuring parameters such as gap degree and sector labels. To enhance the readability of the sector labels within the chord diagram, we customized them by wrapping longer labels and adjusting font sizes as necessary. Additionally, for improved clarity, we implemented filters to include only the most significant molecules, considering criteria like their appearance in multiple pathways or highest significance values. The final output was a chord diagram that effectively portrayed the associations between the top canonical pathways and enriched genes, achieved through the capabilities of the circlize package.

## Acknowledgements

We acknowledge support for bioinformatics analysis from the Harvard NeuroDiscovery Center and the Harvard Chan Bioinformatics Core, as well as funding support from R01 DK077195 (RMA), T32 DK060442 (RMA), R01 DK12 6 3 (RSL, RMA) and the Children’s Urological Foundation.

## Disclosure and competing interests statement

The authors state that no competing interests exist and there are no relevant financial or non-financial interests to disclose.

## Availability of published material, data and software

RNA sequencing data have been deposited in the European Nucleotide Archive (accession number PRJEB67472). The mass spectrometry proteomics data have been deposited to the ProteomeXchange Consortium via the PRIDE [1] partner repository with the dataset identifier PXD046096.

## Figure legends for Supplementary Figures

**Figure Supplementary 1:**
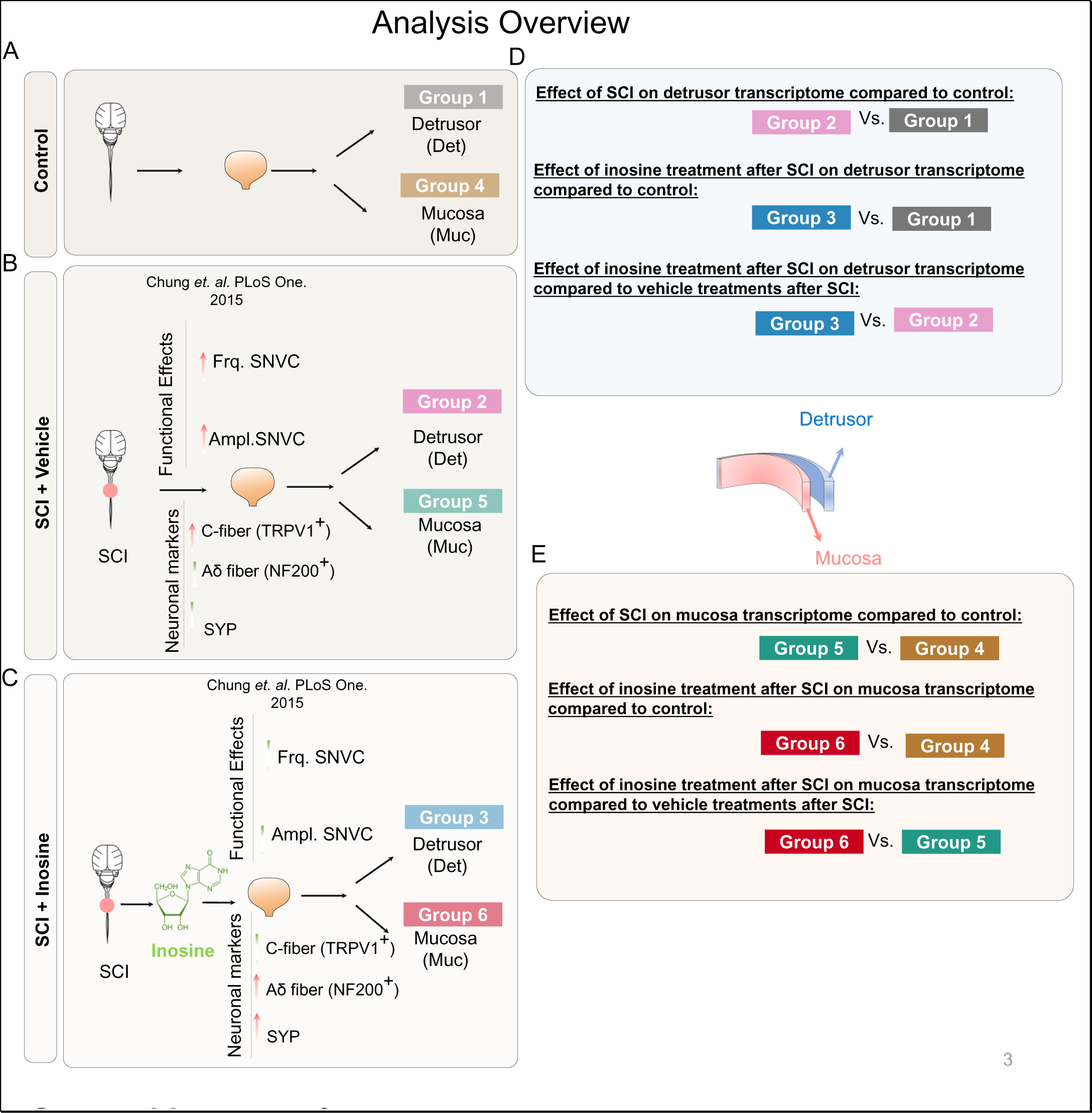
Experimental Group Visualization and Transcriptome Comparisons. (A) Visualization of 6 Distinct Experimental Groups in the RNA Sequencing Experiment of this Study: Group 1 represents detrusor samples from the control uninjured bladder, while Group 4 corresponds to mucosa samples from the same control uninjured bladder. (B) Group 2 comprises detrusor samples from the spinal cord-injured (SCI) bladder treated with a vehicle, and Group 5 consists of mucosa samples from the same SCI bladder treated with the same vehicle. This panel also presents results from our previous studies, highlighting decreased levels of synaptophysin and the Ad fiber marker NF200 in the bladder wall following SCI compared to non-injured controls. (C) Group 3 includes detrusor samples from the SCI bladder treated with inosine, and Group 6 consists of mucosa samples from the same SCI bladder treated with inosine. This panel also showcases the mitigating effect of inosine on neurogenic detrusor overactivity in spinal cord-injured rats, as observed in our previous studies. (D) Three distinct comparison groups within the detrusor: the effect of SCI on the detrusor transcriptome compared to control. the impact of inosine treatment post-SCI on the detrusor transcriptome compared to control. the effect of inosine treatment post-SCI on the detrusor transcriptome compared to vehicle treatment post-SCI. (E) Three distinct comparison groups within the mucosa: effect of SCI on the mucosa transcriptome compared to control. the influence of inosine treatment post-SCI on the mucosa transcriptome compared to control. the effect of inosine treatment post-SCI on the mucosa transcriptome compared to vehicle treatment post-SCI.

**Figure Supplementary 2.**
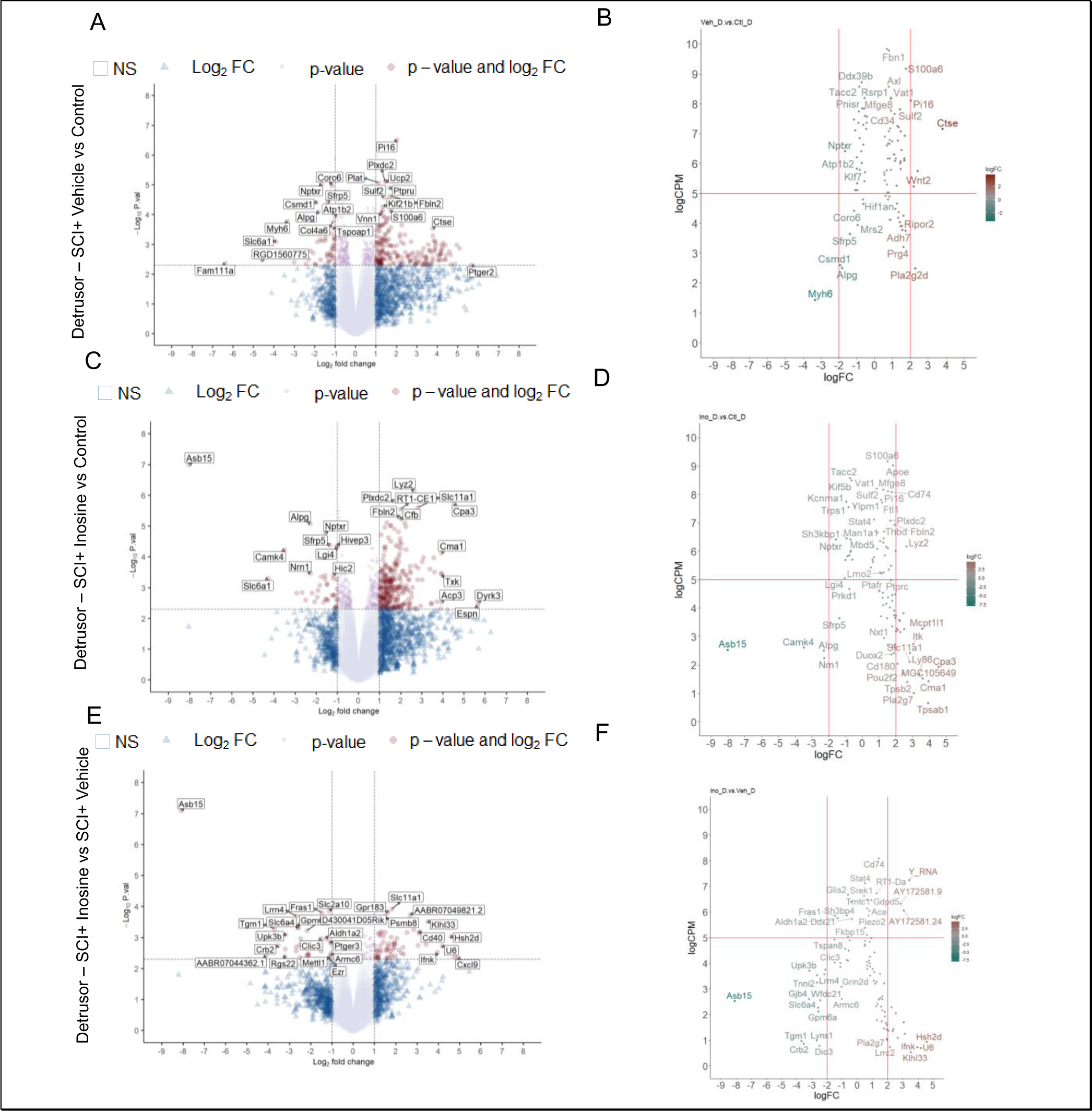
Differentially expressed genes (DEGs) in detrusor. (A) Volcano plot of DEGs in detrusor of SCI treated with vehicle versus control. (B) Scatterplot of DEGs in detrusor of SCI treated with vehicle versus control. Scatterplot shows logFC (fold change) on the x-axis and logCPM (counts per million) on the y-axis, where each point represents a gene. The color of each point is determined by its logFC value, with a gradient ranging from green (negative logFC values, downregulated genes) to red (positive logFC values, upregulated genes), with a gray midpoint representing genes with a logFC value of zero. The red horizontal line represents the cutoff for statistical significance (PValue < 0.05), while the red vertical lines indicate a logFC of −2 and 2, representing the thresholds for down- and upregulation, respectively.(C) Volcano plot of DEGs in detrusor of SCI treated with inosine versus control. (D) Scatterplot of DEGs in detrusor of SCI treated with inosine versus control. X-axis represents the logarithmic fold change and y-axis shows the logarithmic count per million (as a measure for level of expression). (E) Volcano plot of DEGs in detrusor of SCI treated with inosine versus SCI treated with vehicle. This volcano plot represents the subtraction of genes shown in panel A and C. (F) Scatterplot of DEGs in detrusor of SCI treated with inosine versus SCI treated with vehicle. X-axis represents the logarithmic fold change and y-axis shows the logarithmic count per million (as a measure for level of expression). This scatterplot represents the subtraction of genes shown in panel B and D.

**Figure Supplementary 3.**
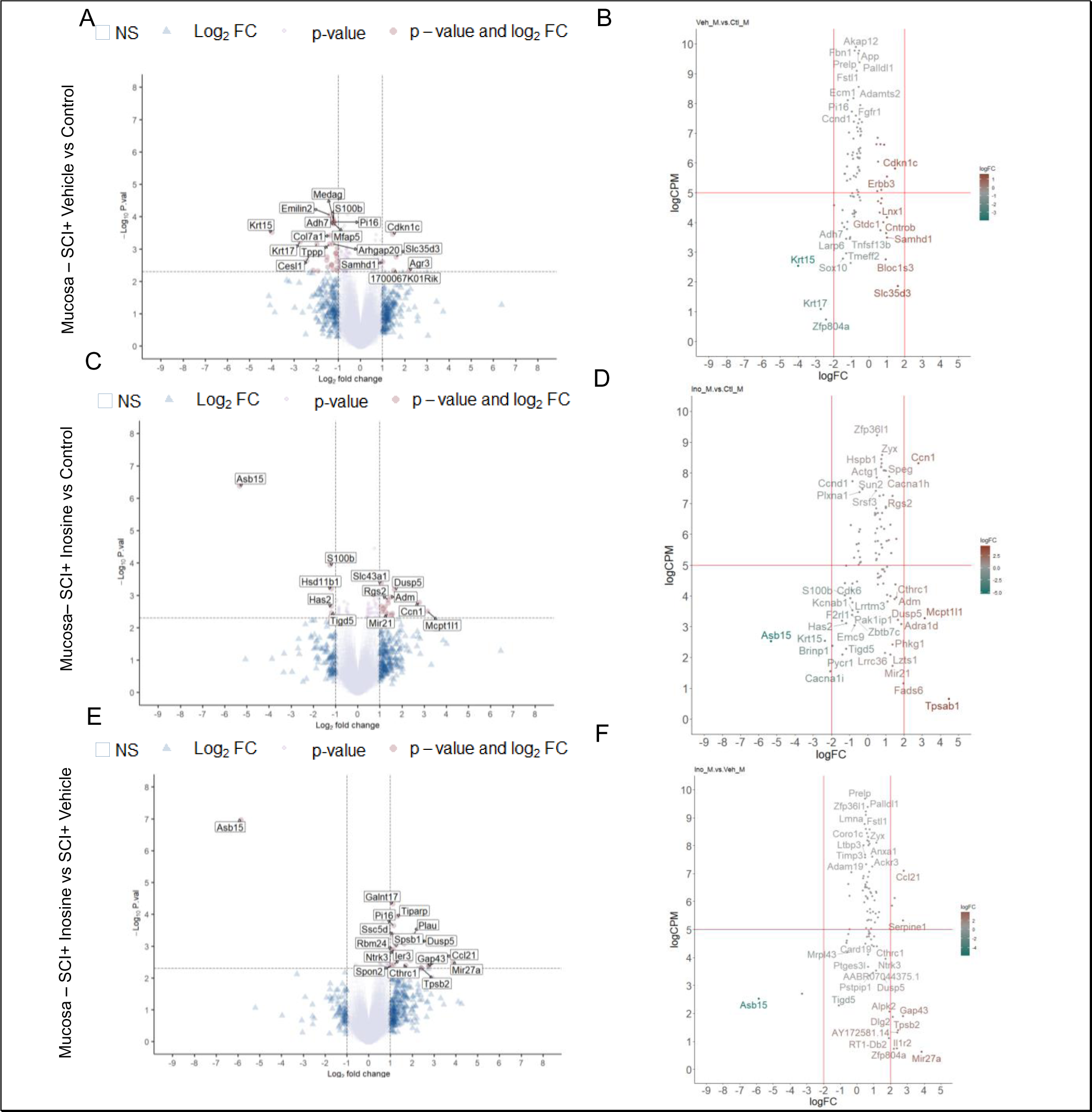
Differentially expressed genes (DEGs) in mucosa. (A) Volcano plot of DEGs in detrusor of SCI treated with vehicle versus control. (B) Scatterplot of DEGs in detrusor of SCI treated with vehicle versus control. X-axis represents the logarithmic fold change and y-axis shows the logarithmic count per million (as a measure for level of expression). (C) Volcano plot of DEGs in detrusor of SCI treated with vehicle versus control. (D) Scatterplot of DEGs in detrusor of SCI treated with inosine versus control. X-axis represents the logarithmic fold change and y-axis shows the logarithmic count per million (as a measure for level of expression). (E) Volcano plot of DEGs in detrusor of SCI treated with inosine versus SCI treated with vehicle. This volcano plot represents the subtraction of genes shown in panel A and C. (F) Scatterplot of DEGs in detrusor of SCI treated with inosine versus SCI treated with vehicle. X-axis represents the logarithmic fold change and y-axis shows the logarithmic count per million (as a measure for level of expression). This scatterplot represents the subtraction of genes shown in panel B and D.

**Figure Supplementary 4.**
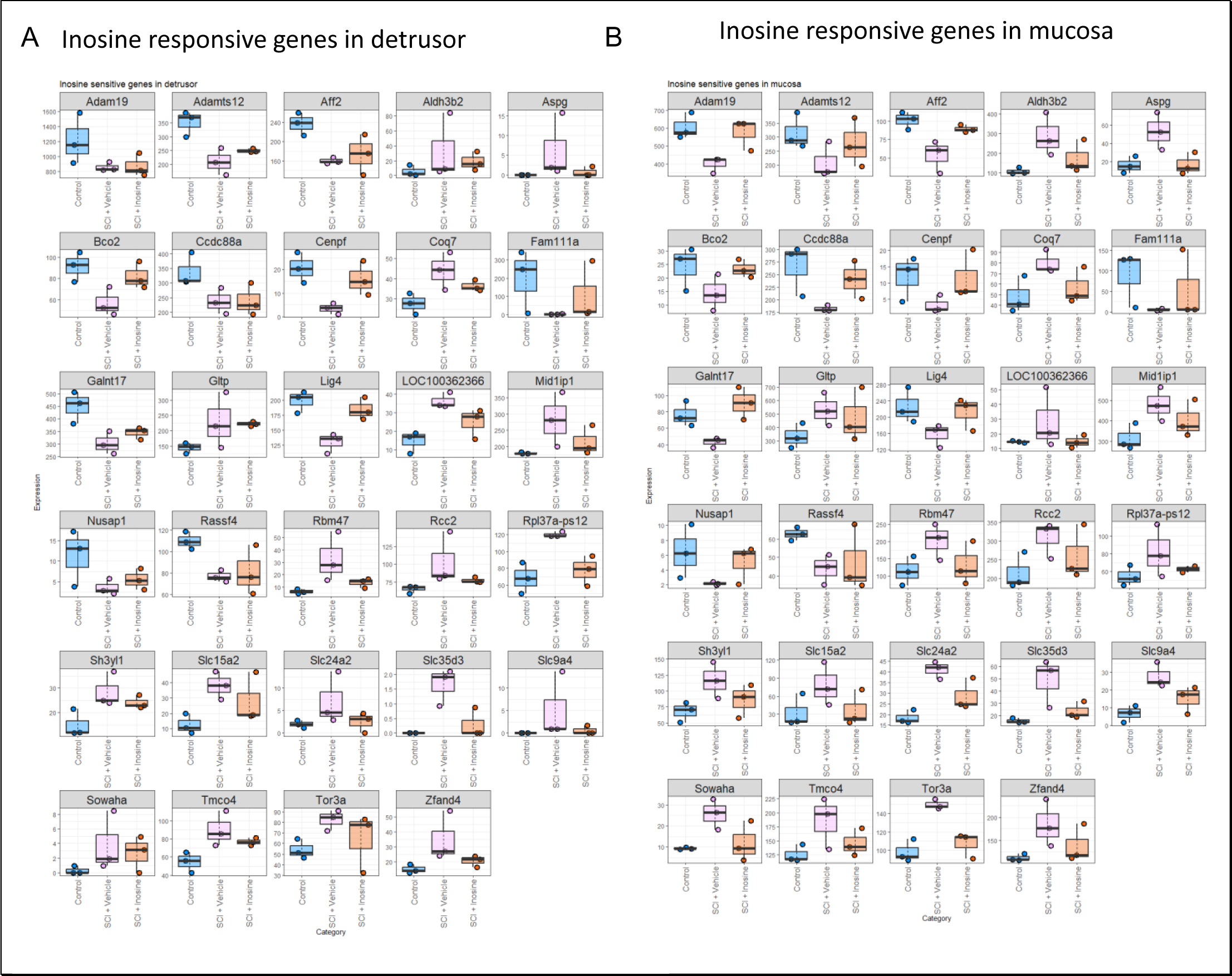
Expression Levels of Inosine-Responsive Genes in Detrusor and Mucosa. (A) Box chart depicting the expression levels of Inosine-responsive genes in the detrusor. The groups include control samples (blue), detrusor samples from spinal cord-injured rats treated with a vehicle (pink), and detrusor samples from spinal cord-injured rats treated with inosine (orange). (B) Box chart illustrating the expression levels of Inosine-responsive genes in the mucosa. The groups consist of control samples, mucosa samples from spinal cord-injured rats treated with a vehicle, and mucosa samples from spinal cord-injured rats treated with inosine.”

**Figure Supplementary 5.**
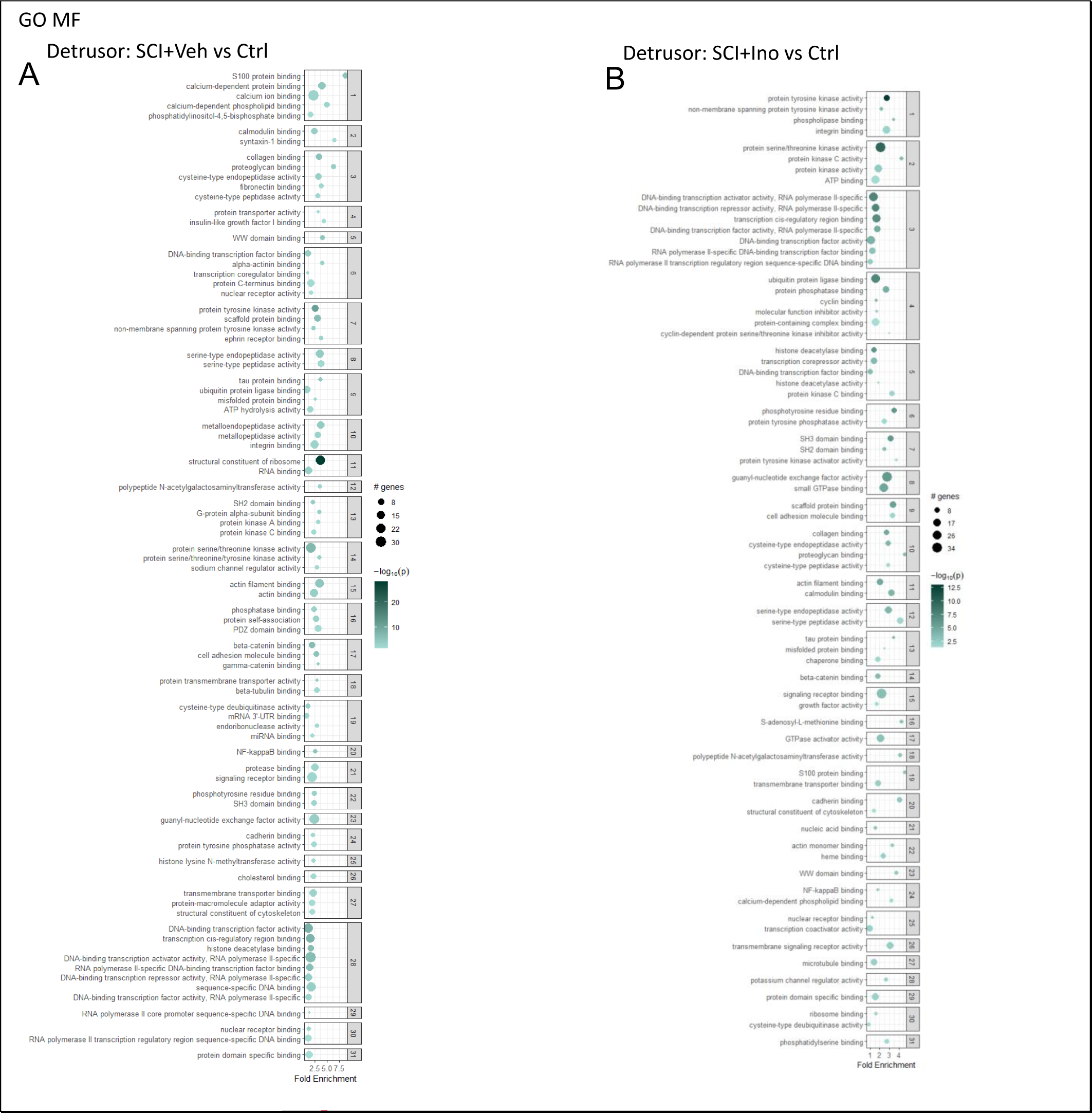
Hierarchically Clustered GO Molecular Functions (MFs) Based on Regulated RNAs in Detrusor of Spinal Cord-Injured (SC) Rats. (A) Hierarchically Clustered GO Molecular Functions (MFs) based on regulated RNAs in the detrusor of SC rats treated with vehicle compared to controls.(B) Hierarchically Clustered GO Molecular Functions (MFs) based on regulated RNAs in the detrusor of SC rats treated with Inosine compared to controls. The intensity or shade of green in the bubbles represents the significance or enrichment of the Gene Ontology (GO) Molecular Function (MF) term. Darker, more intense green indicates higher significance or stronger enrichment of the term. The size of the bubbles corresponds to the number of genes associated with the particular GO MF term. Larger bubbles indicate that a greater number of genes in your dataset are associated with that specific MF term. To simplify the interpretation of our enrichment analysis results, we implemented a clustering approach to group related terms. This approach led to the creation of an enrichment bubble chart that visually presents the clustered terms. We utilized hierarchical clustering, employing the 1 - kappa statistic as the distance metric. The optimal number of clusters was determined by maximizing the average silhouette width, ensuring meaningful groupings. Subsequently, we selected representative terms from each cluster based on their lowest p-values

**Figure Supplementary 6.**
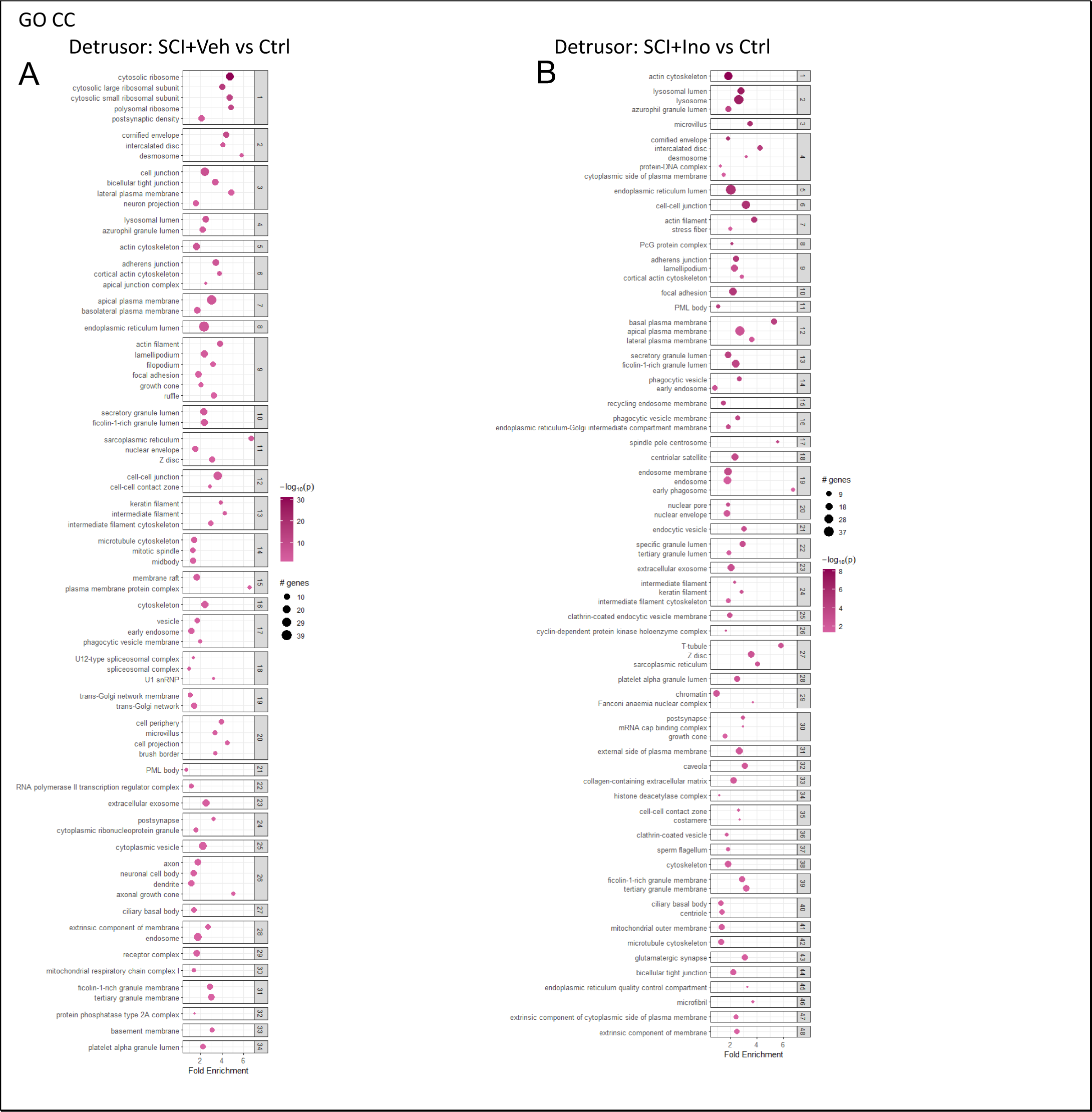
Hierarchically Clustered GO Cellular Components (CCs) Based on Regulated RNAs in Detrusor of Spinal Cord-Injured (SC) Rats. (A) Hierarchically Clustered GO Cellular Components (CCs) based on regulated RNAs in the detrusor of SC rats treated with vehicle compared to controls.(B) Hierarchically Clustered GO Molecular Functions (MFs) based on regulated RNAs in the detrusor of SC rats treated with Inosine compared to controls.The intensity or shade of green in the bubbles represents the significance or enrichment of the Gene Ontology (GO) Cellular Components (CCs) term. Darker, more intense pink indicates higher significance or stronger enrichment of the term. The size of the bubbles corresponds to the number of genes associated with the particular GO CC term. Larger bubbles indicate that a greater number of genes in your dataset are associated with that specific MF term. To simplify the interpretation of our enrichment analysis results, we implemented a clustering approach to group related terms. This approach led to the creation of an enrichment bubble chart that visually presents the clustered terms. We utilized hierarchical clustering, employing the 1 - kappa statistic as the distance metric. The optimal number of clusters was determined by maximizing the average silhouette width, ensuring meaningful groupings. Subsequently, we selected representative terms from each cluster based on their lowest p-values

**Figure Supplementary 7.**
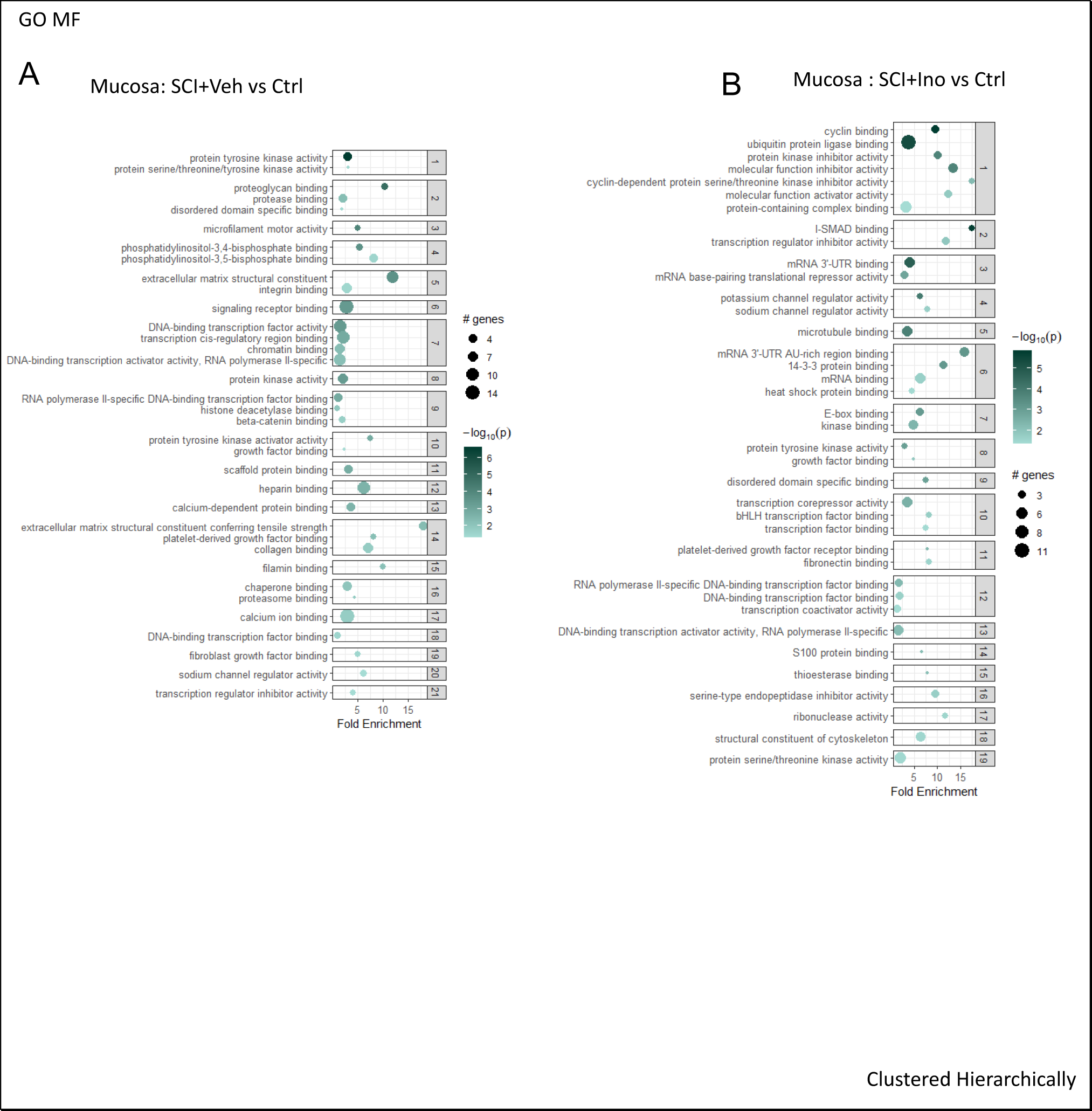
Hierarchically Clustered GO Molecular Functions (MFs) Based on Regulated RNAs in Mucosa of Spinal Cord-Injured (SC) Rats. (A) Hierarchically Clustered GO Molecular Functions (MFs) based on regulated RNAs in the mucosa of SC rats treated with vehicle compared to controls.(B) Hierarchically Clustered GO Molecular Functions (MFs) based on regulated RNAs in the mucosa of SC rats treated with Inosine compared to controls. The intensity or shade of green in the bubbles represents the significance or enrichment of the Gene Ontology (GO) Molecular Function (MF) term. Darker, more intense green indicates higher significance or stronger enrichment of the term. The size of the bubbles corresponds to the number of genes associated with the particular GO MF term. Larger bubbles indicate that a greater number of genes in your dataset are associated with that specific MF term. To simplify the interpretation of our enrichment analysis results, we implemented a clustering approach to group related terms. This approach led to the creation of an enrichment bubble chart that visually presents the clustered terms. We utilized hierarchical clustering, employing the 1 - kappa statistic as the distance metric. The optimal number of clusters was determined by maximizing the average silhouette width, ensuring meaningful groupings. Subsequently, we selected representative terms from each cluster based on their lowest p-values

**Figure Supplementary 8.**
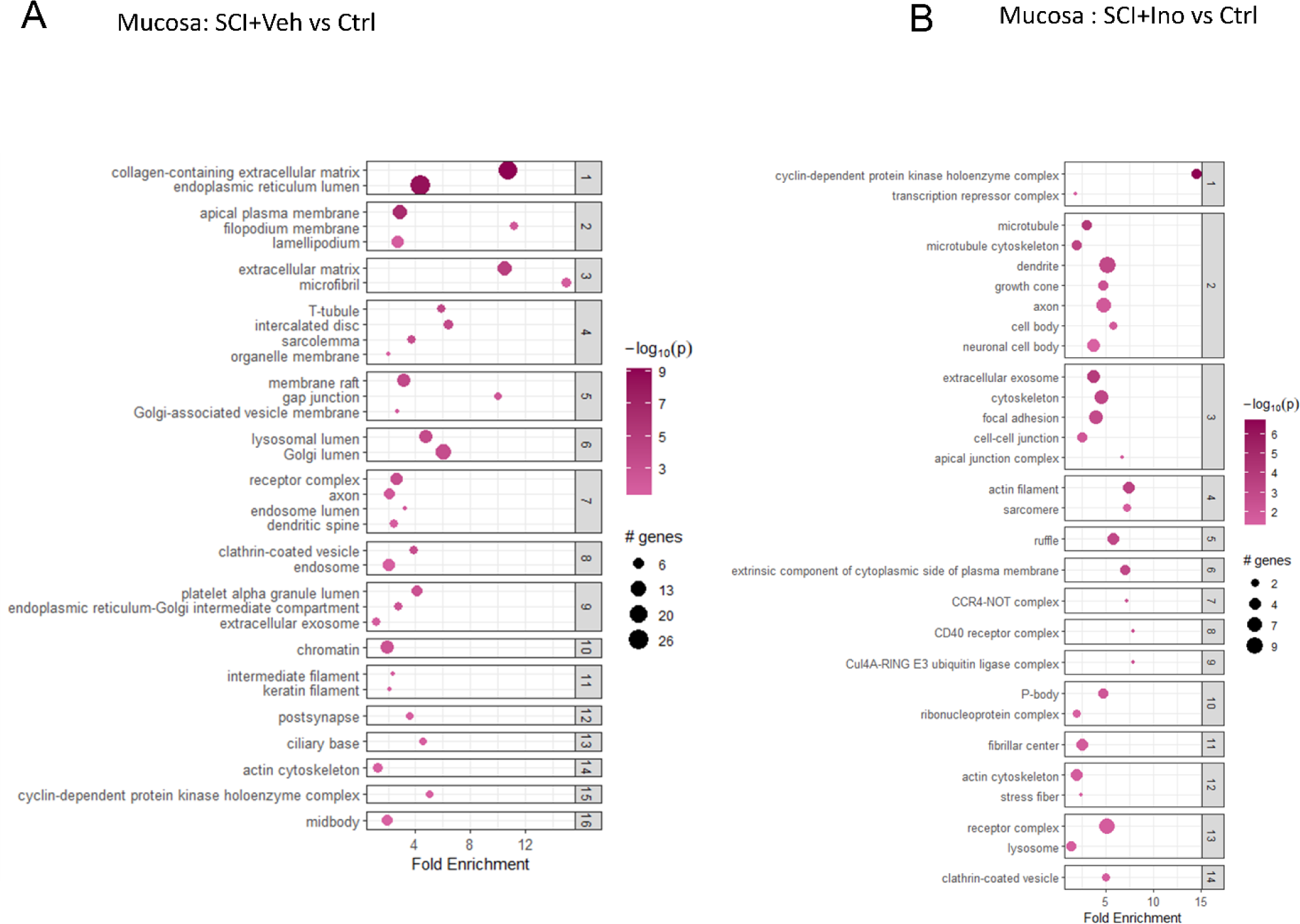
Hierarchically Clustered GO Cellular Components (CCs) Based on Regulated RNAs in Mucosa of Spinal Cord-Injured (SC) Rats. (A) Hierarchically Clustered GO Cellular Components (CCs) based on regulated RNAs in the mucosa of SC rats treated with vehicle compared to controls.(B) Hierarchically Clustered GO Cellular Components (CC) based on regulated RNAs in the mucosa of SC rats treated with Inosine compared to controls. The intensity or shade of green in the bubbles represents the significance or enrichment of the Gene Ontology (GO) Cellular Components (CCs) term. Darker, more intense pink indicates higher significance or stronger enrichment of the term. The size of the bubbles corresponds to the number of genes associated with the particular GO CC term. Larger bubbles indicate that a greater number of genes in your dataset are associated with that specific MF term. To simplify the interpretation of our enrichment analysis results, we implemented a clustering approach to group related terms. This approach led to the creation of an enrichment bubble chart that visually presents the clustered terms. We utilized hierarchical clustering, employing the 1 - kappa statistic as the distance metric. The optimal number of clusters was determined by maximizing the average silhouette width, ensuring meaningful groupings. Subsequently, we selected representative terms from each cluster based on their lowest p-values

**Figure Supplementary 9.**
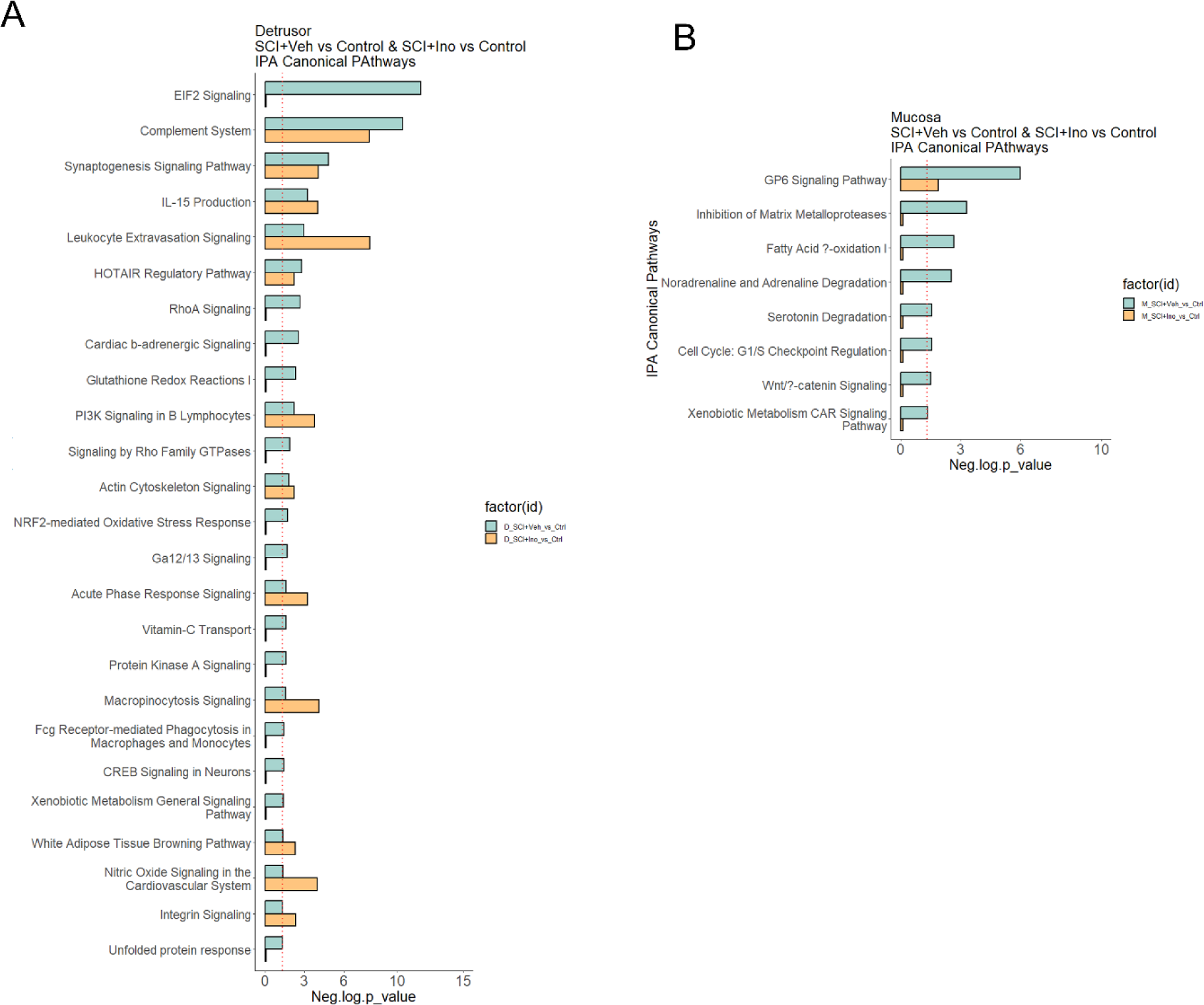
Effect of inosine to preserve the regulated canonical pathways inferred by IPA. Protective Impact of Inosine on Regulated Canonical Pathways in Detrusor (A) and Mucosa (B) as Inferred by IPA. In the bar charts, yellow represents the SCI group treated with inosine compared to the control, while green corresponds to the SCI group treated with a vehicle compared to the control. Thresholds for this analysis were set at Pathways -log10(p-value) cut-off ≤ 1.30 and Path ay z- score cut-off ≥ |1|.

**Figure Supplementary 10.**
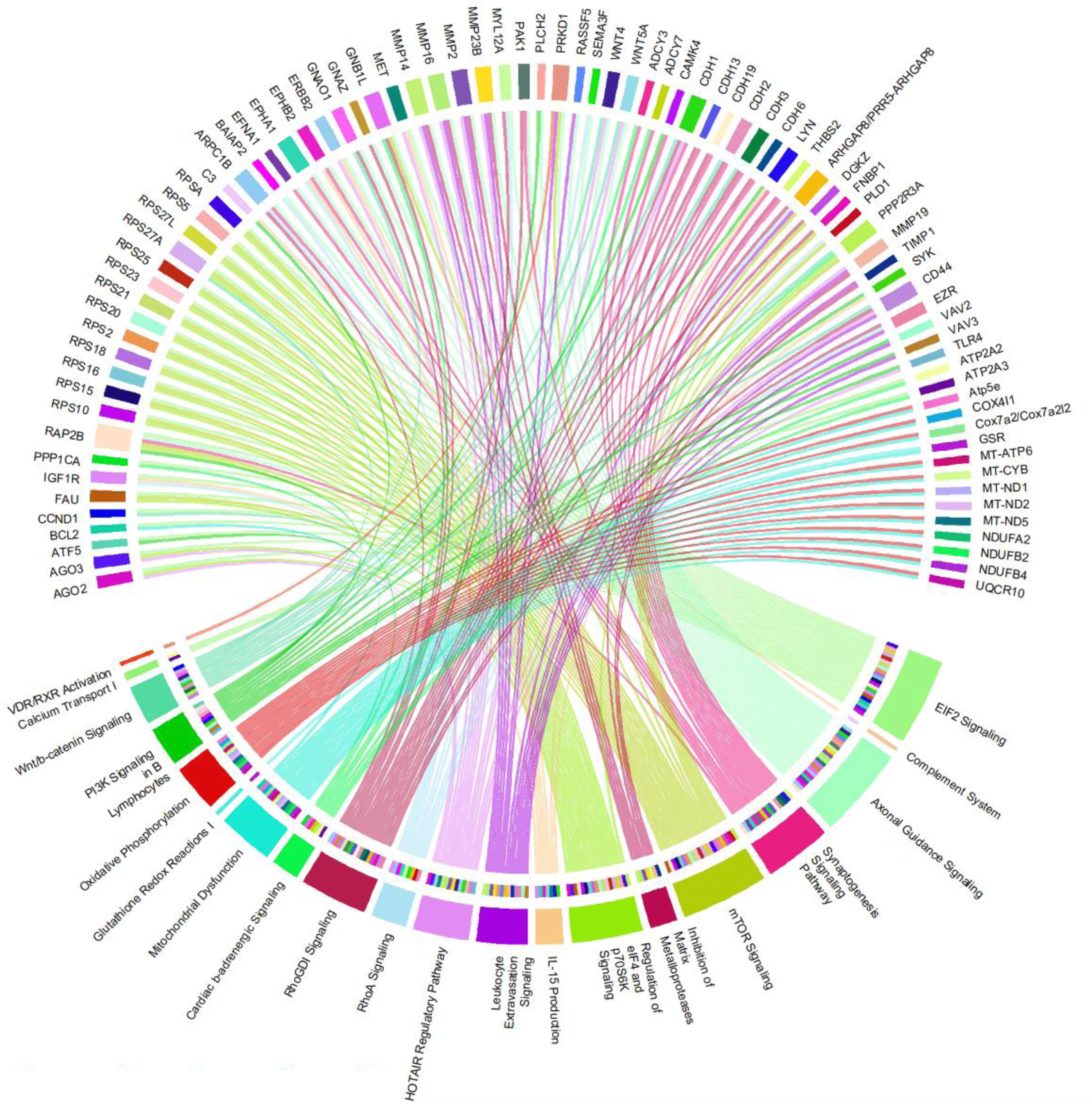
Circos plot of top 20 pathways (based on pvalue) and enriched genes in detrusor of SCI rats treated with vehicle. The plot is a circular diagram that consists of several components: Sectors: The outer circle is divided into sectors, each representing a pathway or a molecule. These sectors are labelled with the names of the pathways and molecules. Links (chords): The lines (chords) connecting the sectors represent relationships between molecules (proteins) and pathways.

**Figure Supplementary 11.**
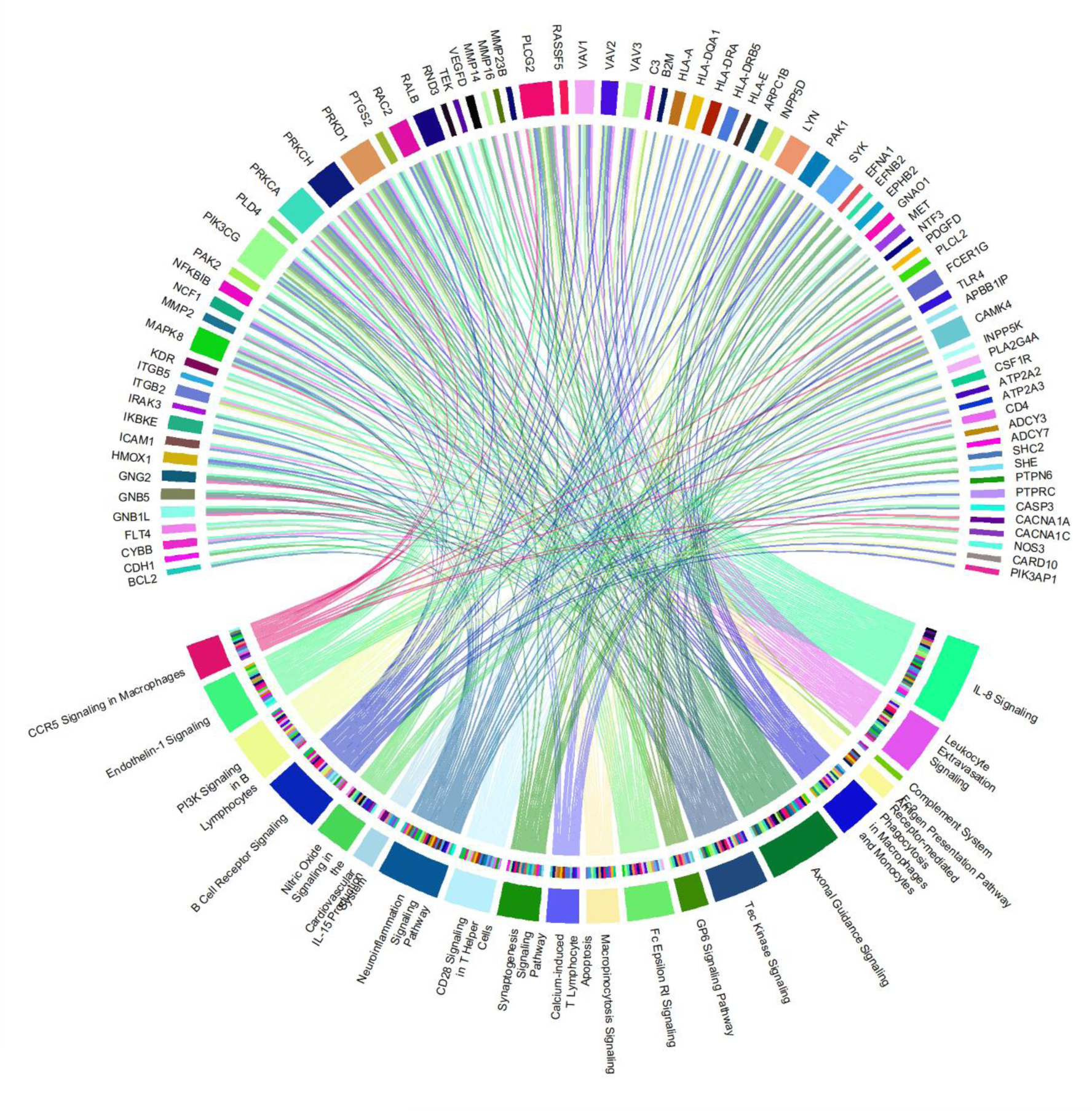
Circos plot of top 20 pathways (based on pvalue) and enriched genes in detrusor of SCI rats treated with inosine. The plot is a circular diagram that consists of several components: Sectors: The outer circle is divided into sectors, each representing a pathway or a molecule. These sectors are labelled with the names of the pathways and molecules. Links (chords): The lines (chords) connecting the sectors represent relationships between molecules (proteins) and pathways.

**Figure Supplementary 12.**
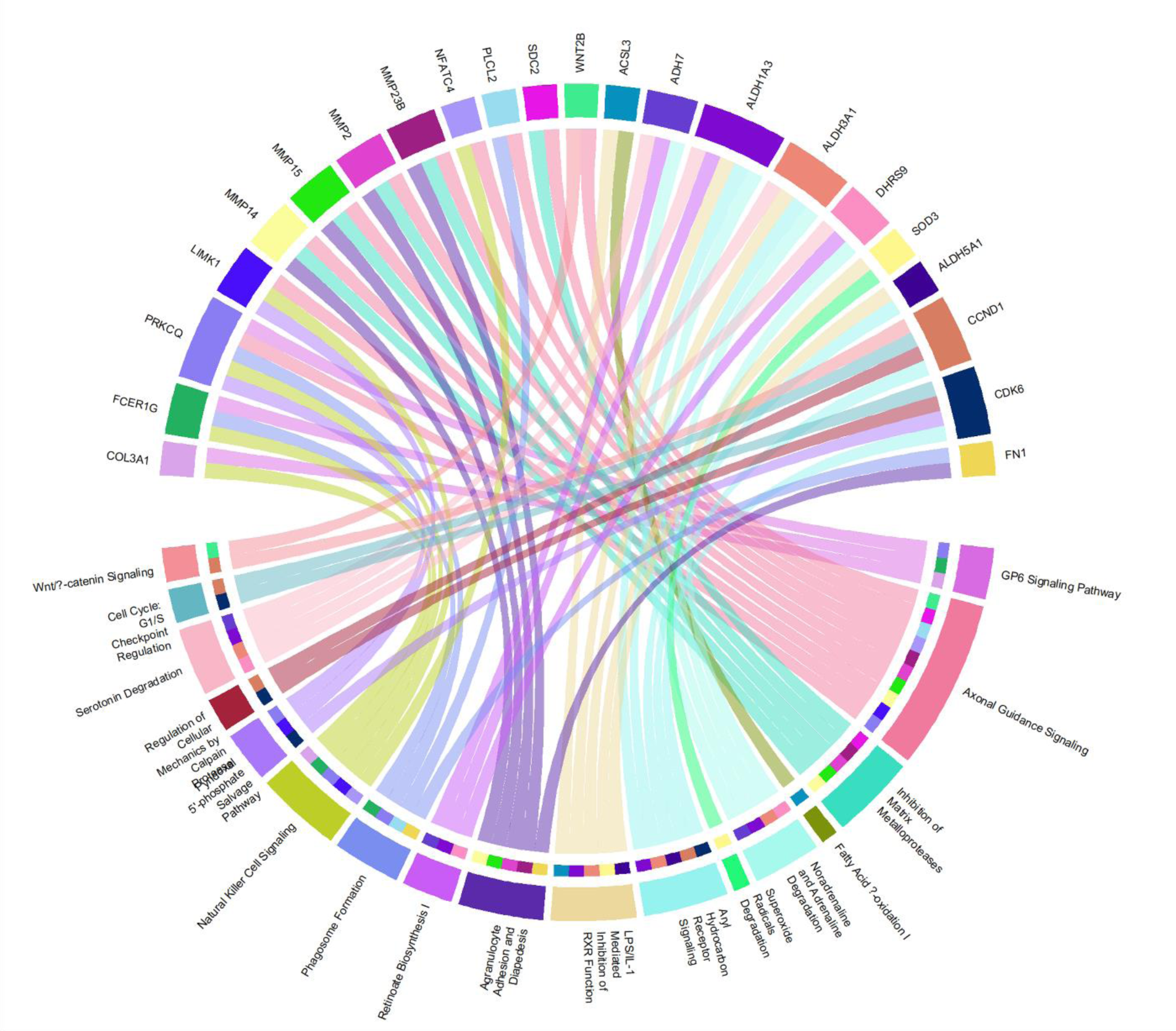
Circos plot of top 20 pathways (based on pvalue) and enriched genes in mucosa of SCI rats treated with vehicle. The plot is a circular diagram that consists of several components: Sectors: The outer circle is divided into sectors, each representing a pathway or a molecule. These sectors are labelled with the names of the pathways and molecules. Links (chords): The lines (chords) connecting the sectors represent relationships between molecules (proteins) and pathways.

**Figure Supplementary 13.**
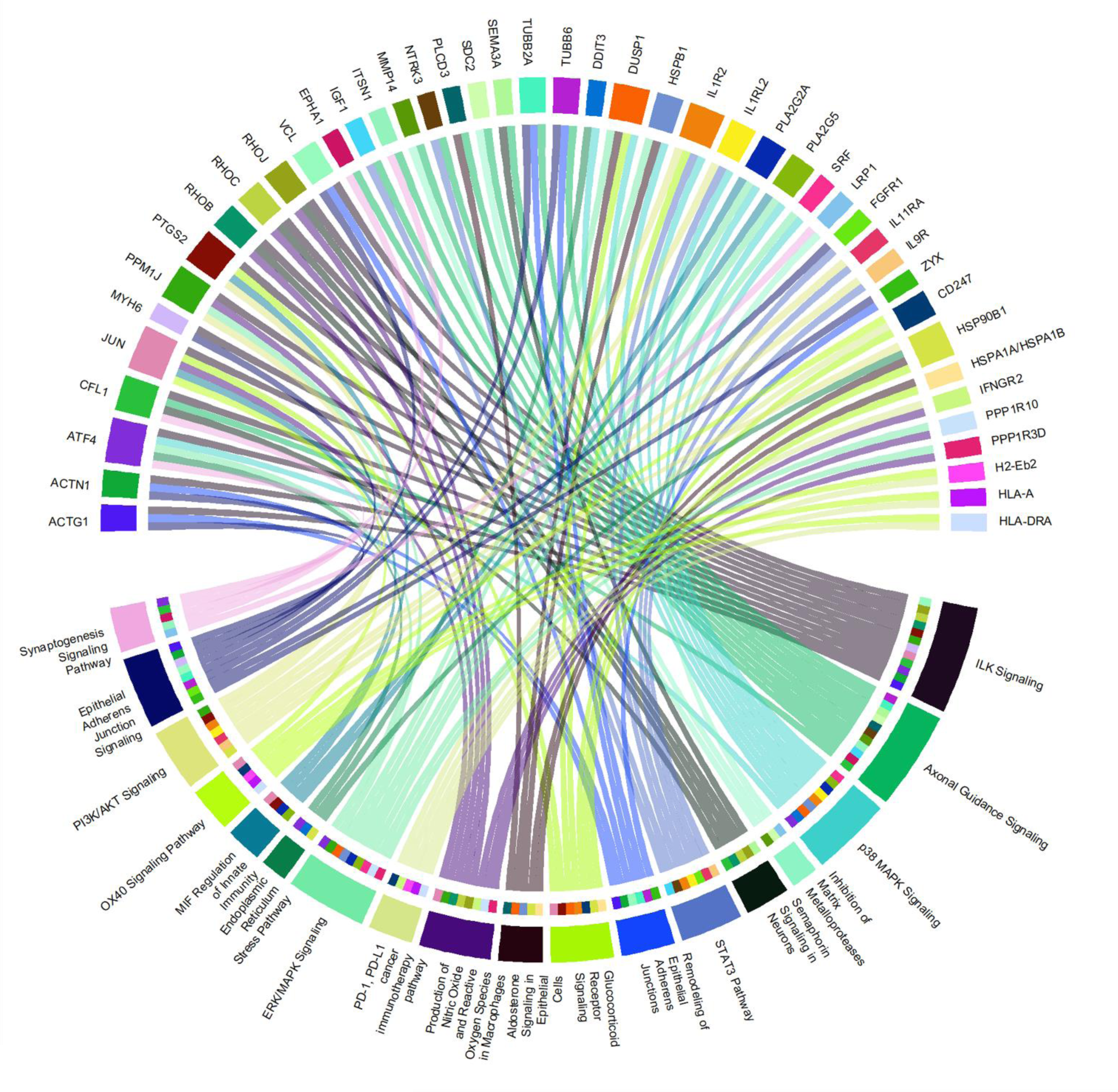
Circos plot of top 20 pathways (based on pvalue) and enriched genes in mucosa of SCI rats treated with inosine. The plot is a circular diagram that consists of several components: Sectors: The outer circle is divided into sectors, each representing a pathway or a molecule. These sectors are labelled with the names of the pathways and molecules. Links (chords): The lines (chords) connecting the sectors represent relationships between molecules (proteins) and pathways.

**Figure Supplementary 14.**
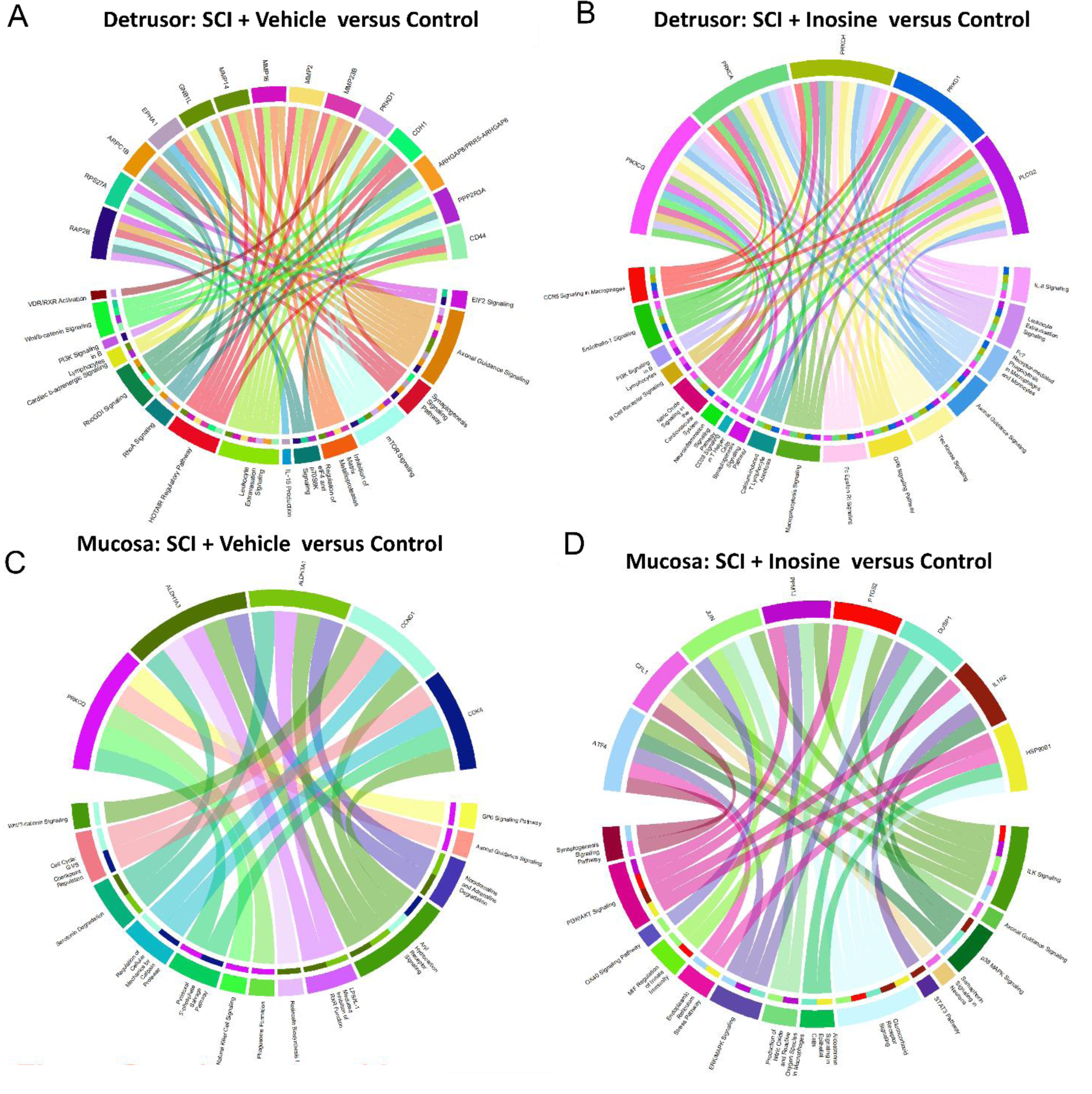
Circos plot of top 20 pathways (based on pvalue) and enriched genes in mucosa of SCI rats treated with vehicle. (A) Circos plot of top recurrent genes in the pathways (reflected in word cloud) in detrusor of SCI rats treated with vehicle versus control.(B) Circos plot of top recurrent genes in the pathways (reflected in word cloud) in detrusor of SCI rats treated with inosine versus control.(C) Circos plot of top recurrent genes in the pathways (reflected in word cloud) in mucosa of SCI rats treated with vehicle versus control. (D) Circos plot of top recurrent genes in the pathways (reflected in word cloud) in mucosa of SCI rats treated with inosine versus control. The plot is a circular diagram that consists of several components: Sectors: The outer circle is divided into sectors, each representing a pathway or a molecule. These sectors are labelled with the names of the pathways and molecules. Links (chords): The lines (chords) connecting the sectors represent relationships between molecules (proteins) and pathways.

**Figure Supplementary 15.**
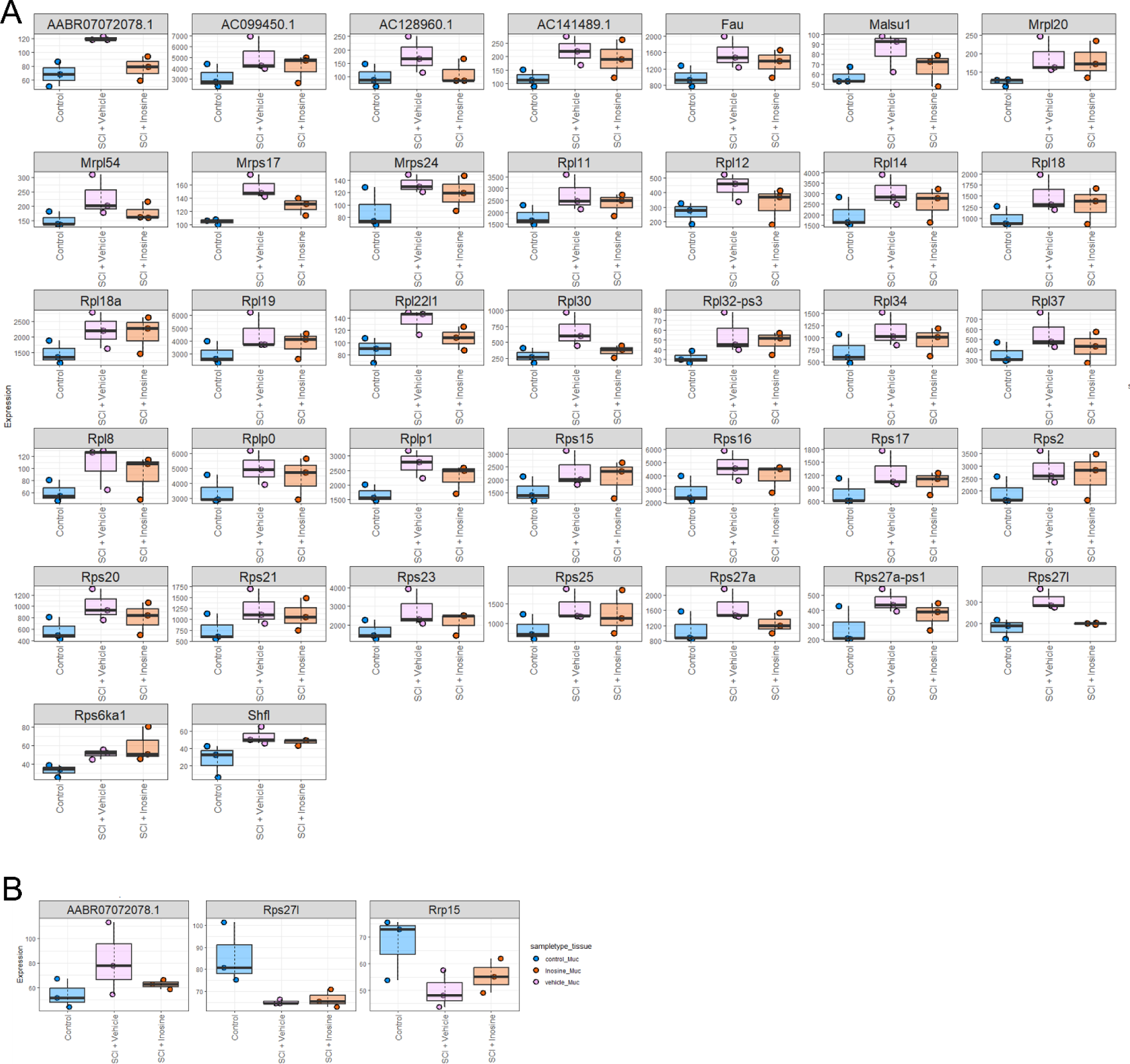
Expression Profiles of Ribosomal Genes in Detrusor and Mucosa of Rats with Spinal Cord Injury. In the data preprocessing workflow, ribosomal genes were selectively identified and extracted from RNA sequencing datasets. The dataset was further refined to include only these ribosomal genes, ensuring that genes with statistical significance (p-value < 0.05) and a substantial change in expression (absolute log-fold change > 0.5) were retained. Expression values across different replicates and conditions (Control, SCI treated with vehicle or inosine in Detrusor (A) or Mucosa (B), were selected.

**Figure Supplementary 16.**
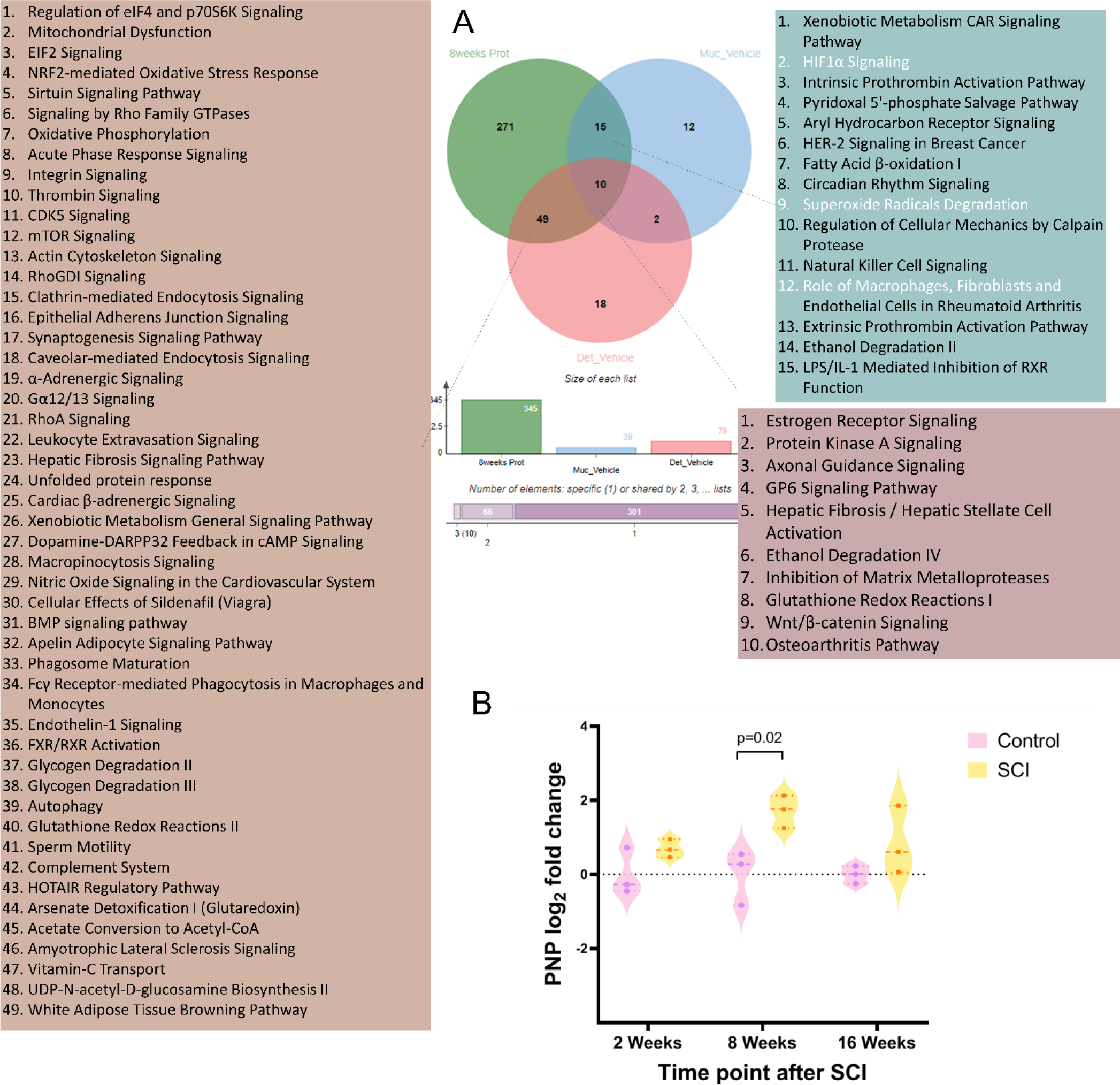
Integration of transcriptomics and proteomics. (A) Illustration of the integration of transcriptomics and proteomics data as a venn diagram that showcases the intersection of pathways between whole bladder proteomics data obtained at 8 weeks post-Spinal Cord Injury (SCI) and RNA sequencing data from both the detrusor and mucosa tissues. This visual representation highlights the commonalities and overlaps in biological pathways between these two types of molecular data. (B) Temporal Expression of Purine Nucleoside Phosphorylase (PNP) Post-Spinal Cord Injury: This violin plot depicts the log2 fold change in PNP expression in rat bladder tissue over a 16-week period after SCI. Data points for both control (pink) and SCI-affected (yellow) groups are displayed at 2-, 8-, and 16-weeks post-injury, with the spread indicating variability within the groups. A statistically significant change (p=0.02) in PNP expression is observed between the control and SCI groups, reflecting the impact of spinal cord injury on this specific protein involved in purine metabolism.

**Table Supplementary 1.**
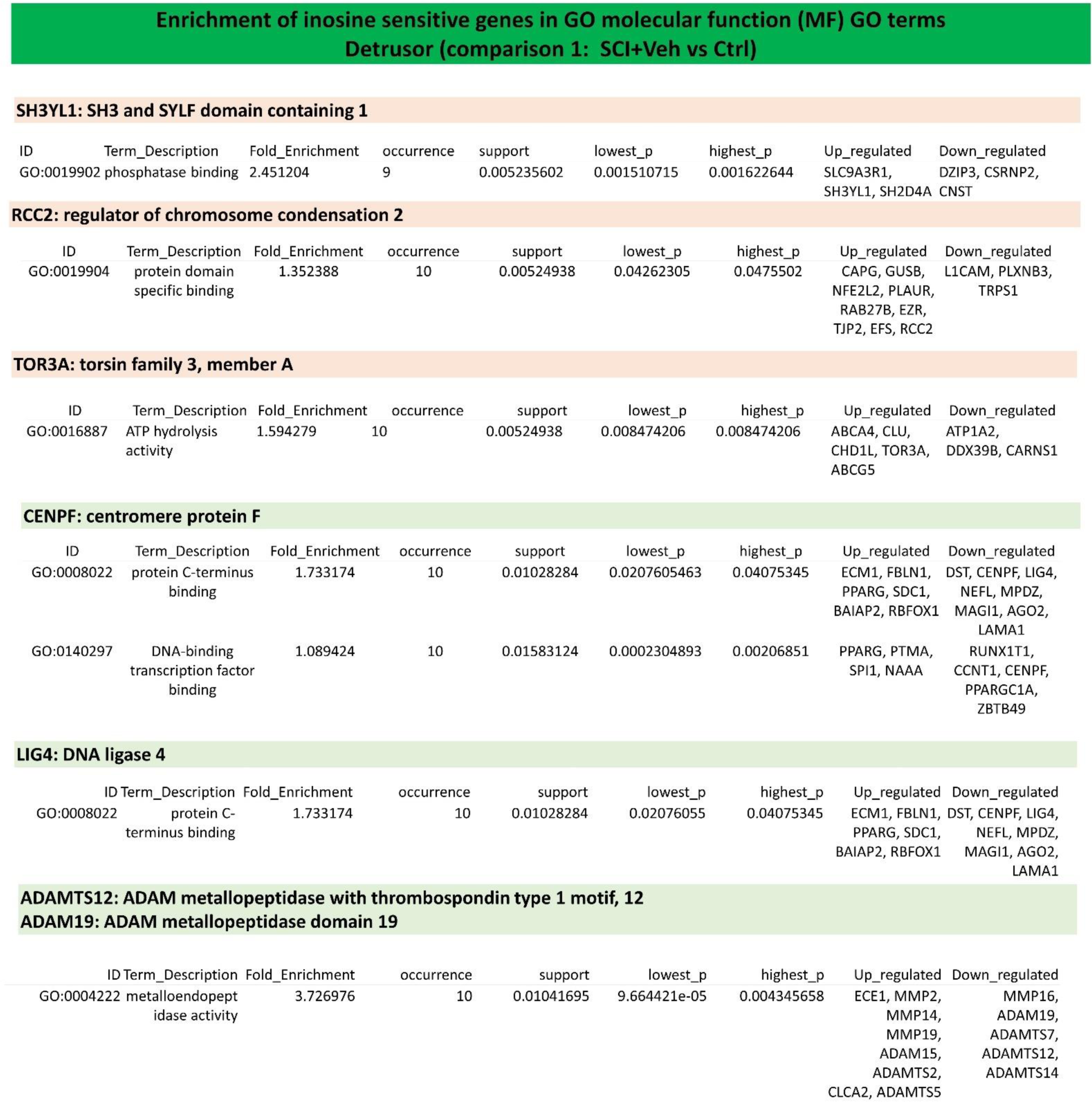
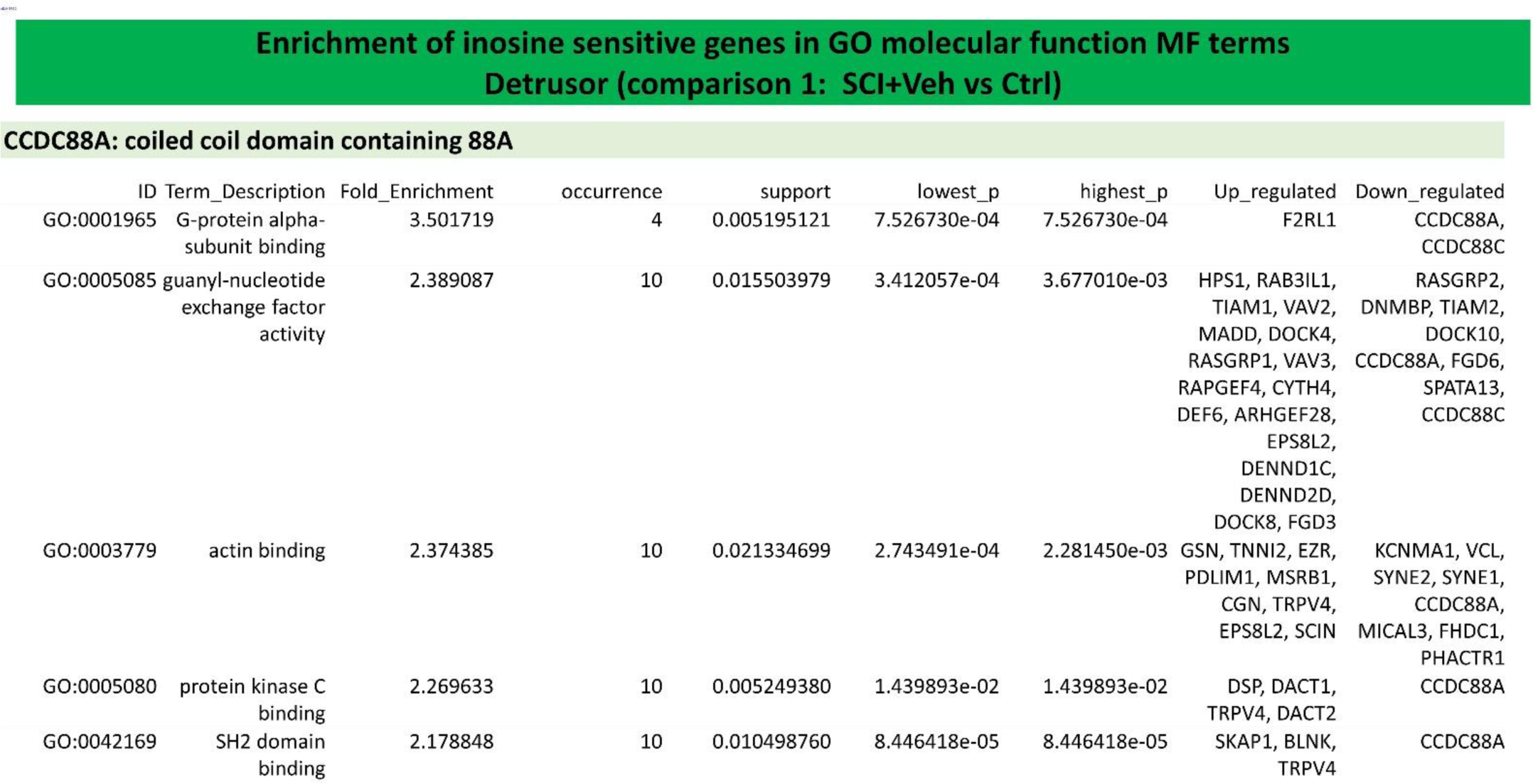
Enrichment of inosine sensitive genes in GO molecular function MF terms Detrusor (SCI+Veh vs Ctrl). Red shad represents genes that were upregulated and inosine treatment has preserved their expression to the control level. Green shad represents genes that were upregulated and inosine treatment has preserved their expression to the control level.

**Table Supplementary 2.**
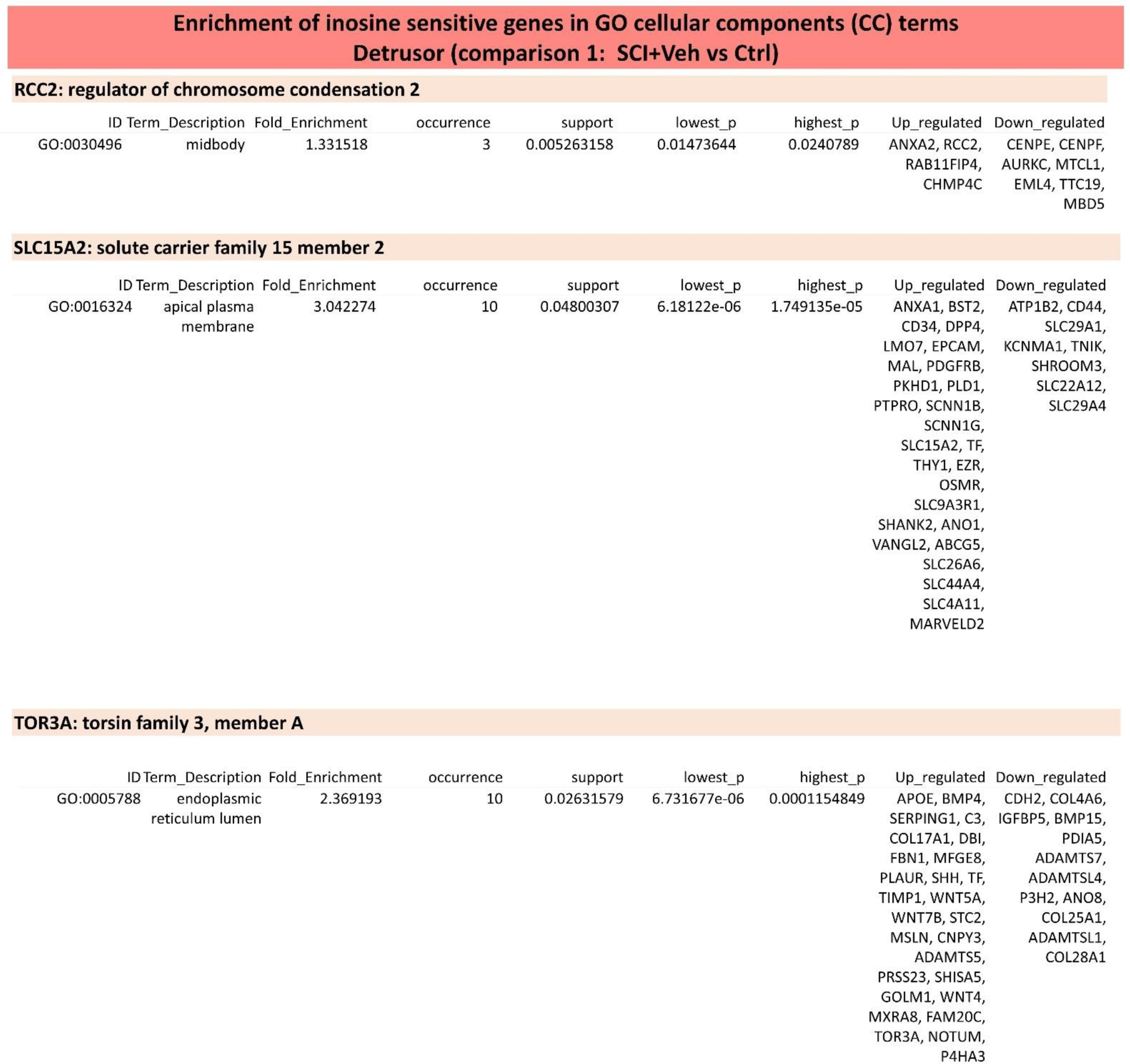
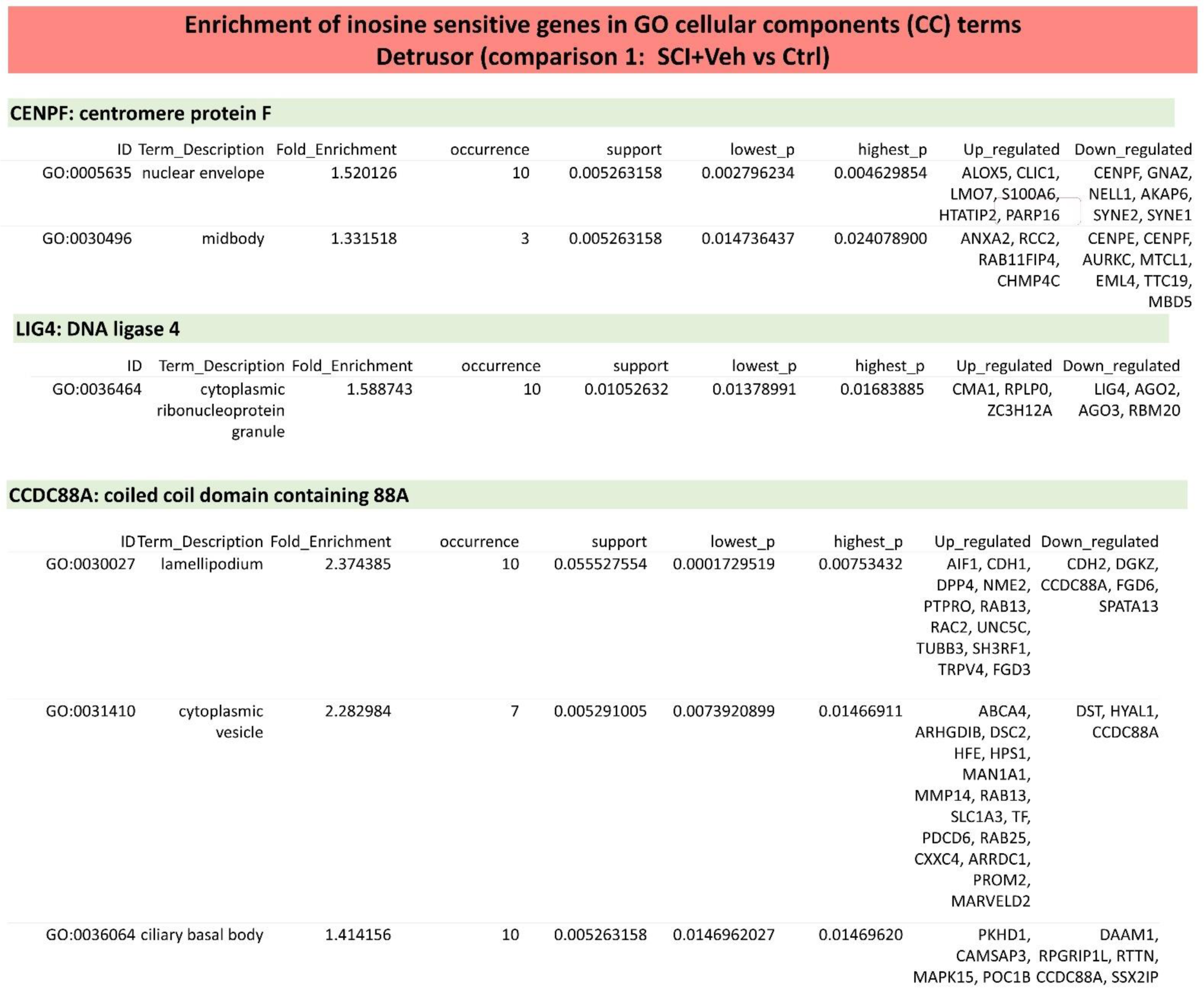
Enrichment of inosine sensitive genes in GO cellular components (CC) terms Detrusor (SCI+Veh vs Ctrl). Red shad represents genes that were upregulated and inosine treatment has preserved their expression to the control level. Green shad represents genes that were upregulated and inosine treatment has preserved their expression to the control level.

**Table Supplementary 3.**
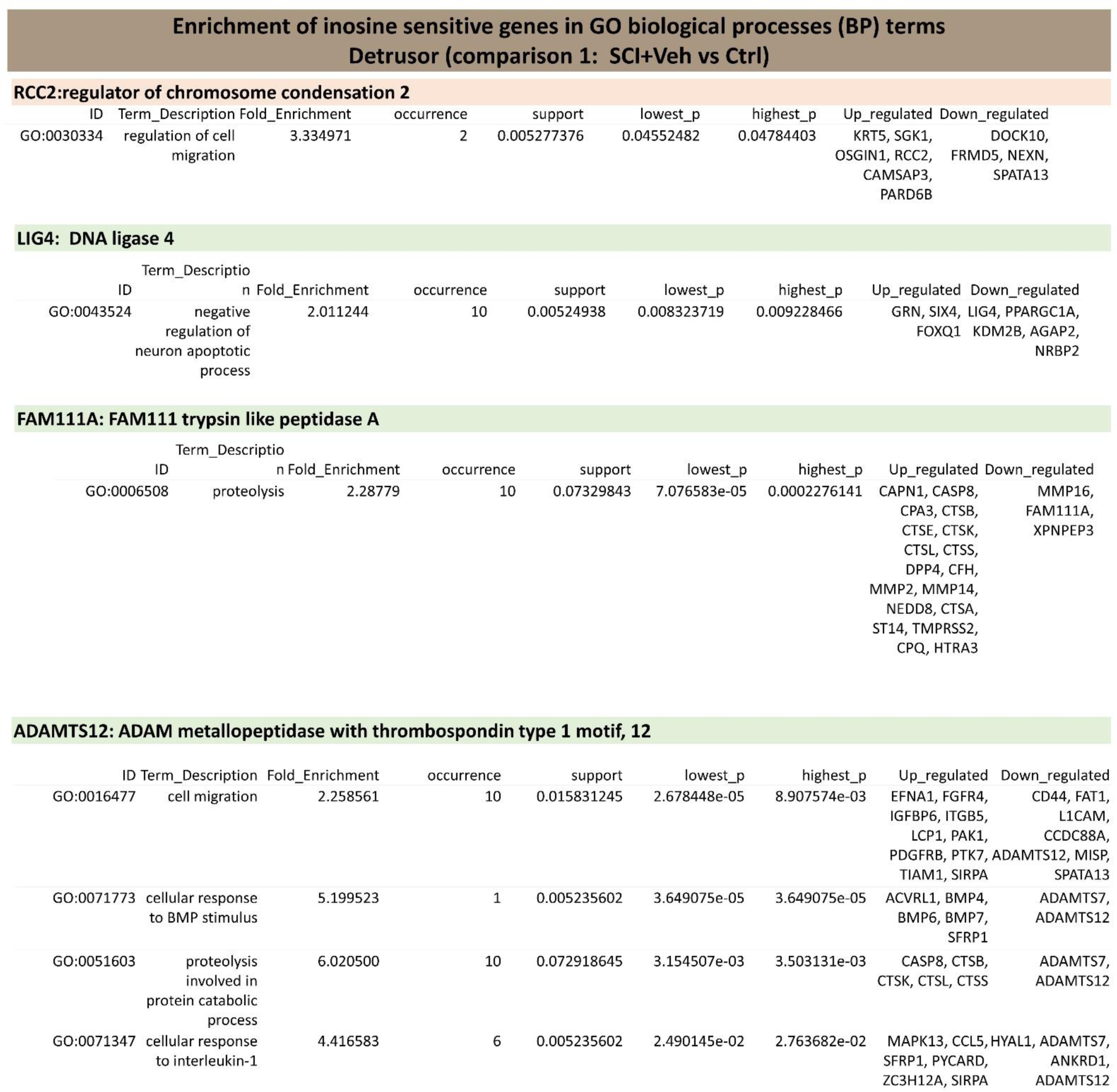
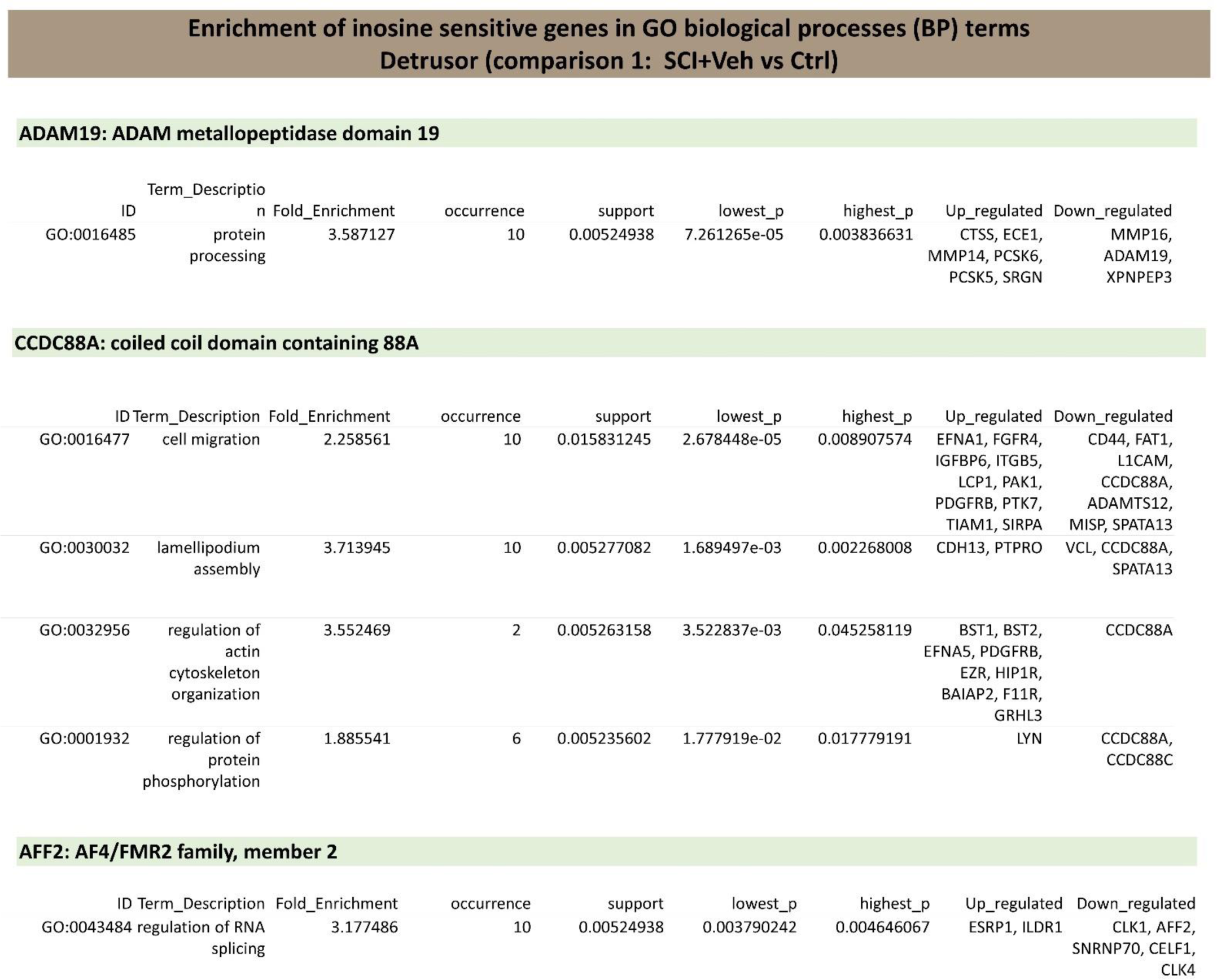
Enrichment of inosine sensitive genes in GO biological processes (BP) terms Detrusor (SCI+Veh vs Ctrl). Red shad represents genes that were upregulated and inosine treatment has preserved their expression to the control level. Green shad represents genes that were upregulated and inosine treatment has preserved their expression to the control level.

**Table Supplementary 4.** Enrichment of inosine sensitive genes in GO molecular function (MF) GO terms Mucosa (SCI+Veh vs Ctrl). Red shad represents genes that were upregulated and inosine treatment has preserved their expression to the control level. Green shad represents genes that were upregulated and inosine treatment has preserved their expression to the control level.

**CENPF: centromere protein F**
| ID | Term_Description | Fold_Enrichment | occurrence | support | lowest_p | highest_p | Up_regulated | Down_regulated |
| --- | --- | --- | --- | --- | --- | --- | --- | --- |
| GO:0140297 | DNA-binding<br>transcription factor<br>binding | 0.9977288 | 10 | 0.005115123 | 0.02956758 | 0.02956758 | HDAC11 | ARNTL, CENPF |

**Table Supplementary 5.**
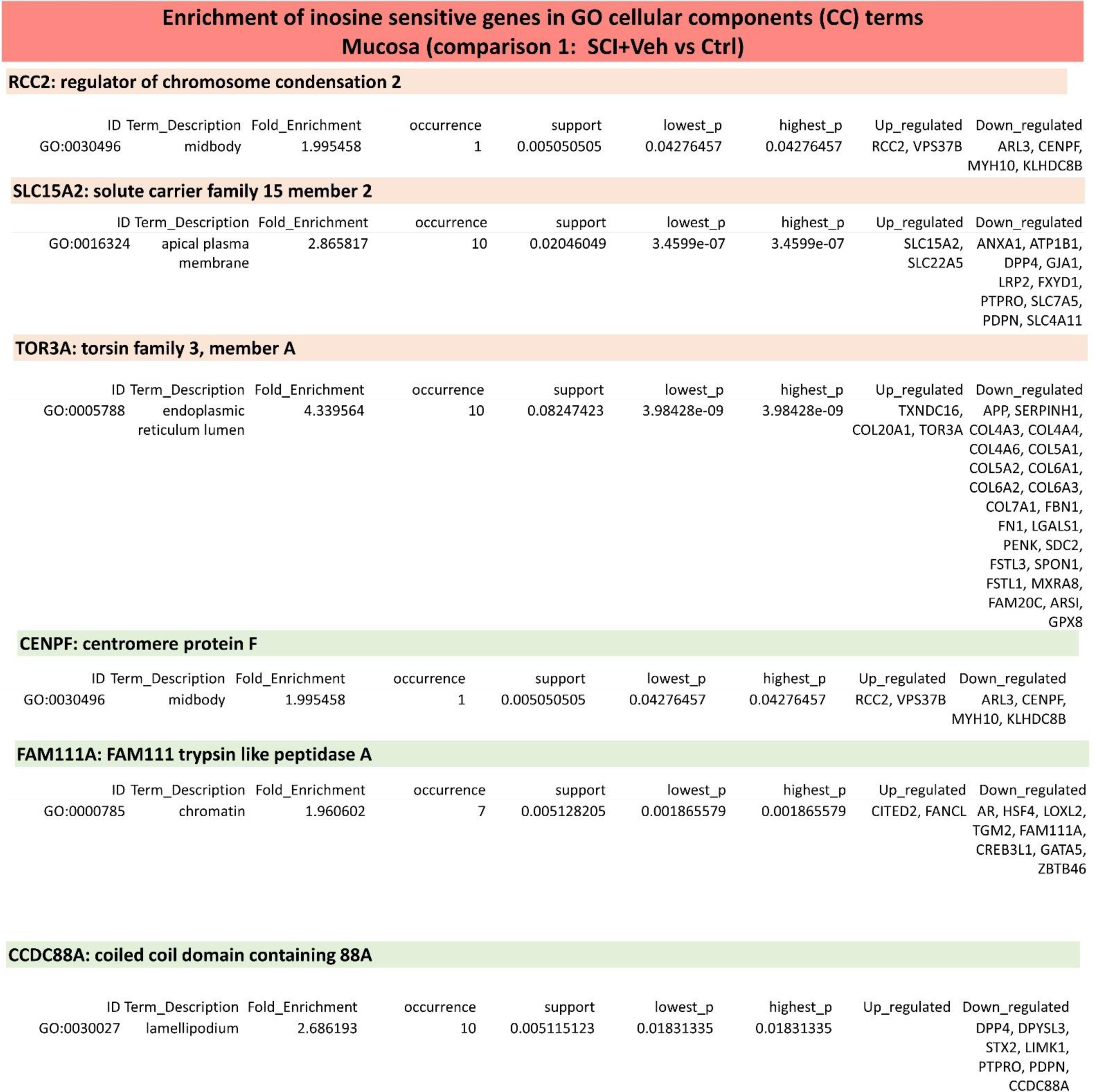
Enrichment of inosine sensitive genes in GO cellular components (CC) terms Mucosa (SCI+Veh vs Ctrl). Red shad represents genes that were upregulated and inosine treatment has preserved their expression to the control level. Green shad represents genes that were upregulated and inosine treatment has preserved their expression to the control level.

**Table Supplementary 6.** Enrichment of inosine sensitive genes in GO biological processes (BP) terms - Mucosa (SCI+Veh vs Ctrl). Red shad represents genes that were upregulated and inosine treatment has preserved their expression to the control level. Green shad represents genes that were upregulated and inosine treatment has preserved their expression to the control level.

**RCC2:regulator of chromosome condensation 2**
| ID | Term_Description | Fold_Enrichment | occurrence | support | lowest_p | highest_p | Up_regulated | Down_regulated |
| --- | --- | --- | --- | --- | --- | --- | --- | --- |
| GO:1900025 | negative regulation of substrate adhesion-dependent cell spreading | 7.482966 | 10 | 0.005102041 | 0.04037285 | 0.04037285 | RCC2 | FBLN1 |

**CCDC88A: coiled coil domain containing 88A**

| ID | Term_Description | Fold_Enrichment | occurrence | support | lowest_p | highest_p | Up_regulated | Down_regulated |
| --- | --- | --- | --- | --- | --- | --- | --- | --- |
| GO:0032148 | activation of protein kinase B activity | 5.612224 | 1 | 0.005102041 | 0.001476416 | 0.001476416 |  | NRG1, NTRK3, CCDC88A |

**AFF2: AF4/FMR2 family, member 2**

| ID | Term_Description | Fold_Enrichment | occurrence | support | lowest_p | highest_p | Up_regulated | Down_regulated |
| --- | --- | --- | --- | --- | --- | --- | --- | --- |
| GO:0010468 | regulation of gene expression | 3.50764 | 10 | 0.01530612 | 0.02487353 | 0.04144516 |  | APP, ATP1B1, CDK6, AFF2, SIRPA |

**NUSAP1: nucleolar and spindle associated protein 1**

| ID | Term_Description | Fold_Enrichment | occurrence | support | lowest_p | highest_p | Up_regulated | Down_regulated |
| --- | --- | --- | --- | --- | --- | --- | --- | --- |
| GO:0000281 | mitotic cytokinesis | 4.380273 | 9 | 0.005102041 | 0.01536256 | 0.020075 | STMN1 | ARL3, MYH10, NUSAP1 |

## Notes

Conflict of interest: The authors declare that no conflict of interest exists.

### Competing Interest Statement

The authors have declared no competing interest.

